# Mutation in mitochondrial chaperone TRAP1 results in male-specific autism

**DOI:** 10.1101/2023.06.02.543381

**Authors:** Małgorzata Rydzanicz, Bozena Kuzniewska, Marta Magnowska, Tomasz Wójtowicz, Ewa Borsuk, Olga Gewartowska, Jakub Gruchota, Anna Hojka, Jacek Miłek, Aleksandra Stawikowska, Patrycja Wardaszka, Izabela Chojnicka, Ludwika Kondrakiewicz, Alicja Puścian, Ewelina Knapska, Andrzej Dziembowski, Rafał Płoski, Magdalena Dziembowska

## Abstract

There is increasing evidence of mitochondrial dysfunction in autism spectrum disorders (ASD), but the causal relationships are unclear. In an ASD patient whose identical twin was unaffected, we identified a postzygotic mosaic mutation p.Q639* in the *TRAP1* gene, which encodes a mitochondrial chaperone of the HSP90 family. Additional screening of 176 unrelated ASD probands revealed an identical *TRAP1* variant in a male patient who had inherited it from a healthy mother. Notably, newly generated knock-in *Trap1* p.Q641* mice display ASD-related behavioral abnormalities exclusively in males. Accordingly, *Trap1* p.Q641* mutation also resulted in sex-specific changes in synaptic plasticity, number of presynaptic mitochondria, and metabolic substrate consumption. Thus, the *TRAP1* p.Q639* mutation is the first example of a monogenic ASD caused by impaired mitochondrial protein homeostasis.

**One-Sentence Summary:** Patient mutation in *TRAP1* causes autism in male mice.

## INTRODUCTION

Autism spectrum disorders (ASD) are a heterogeneous group of early-onset neurodevelopmental disorders with an estimated prevalence of ∼1% in the general population and an average male-to-female ratio of 4-5:1 (*1*). ASD has a complex multifactorial etiology with a strong genetic contribution (*2*). Thanks to the development of large-scale whole-exome studies, many genes associated with ASD have been identified (*3, 4*). Consequently, thousands of gene variants and mutations related to autism have been reported, and it is challenging to identify those most relevant to disease. *De novo* mutations (*5, 6*) and postzygotic mosaic variants (PZMVs) (*7-10*) also contribute to ASD incidence.

Most known human mutations related to autism have been identified in genes encoding synaptic proteins such as neuroligins, neurexins, synapsin 1, Shank proteins, and neurotransmitter receptors (*11-13*). For some of these genes, mouse models with the particular mutations have been generated and showed ASD-like behaviors and synaptic dysfunctions (*14-17*). Analysis of these animal models has supported the concept that the pathophysiology of ASD is related to impaired synaptic function. Synapses are the neuronal regions with the highest energy consumption, making mitochondria particularly essential for synaptic functions. Accordingly, there is growing observational evidence of mitochondrial disturbance in autism spectrum disorders (ASD), but causality of mitochondrial dysfunction in ASD remains to be determined.

Monozygotic twins (MZT), which arise from a single zygote and are considered physically and genetically identical, have been a valuable tool in understanding the extent of the genetic component of many diseases, including autism, for decades. Notably, any genetic differences between MZTs are expected to be postzygotic; thus, disease-discordant MZTs represent an exceptional opportunity to study direct genotype-phenotype correlations. Here, in an ASD-discordant MZT, we identified a p.Q639* nonsense mutation in the *TRAP1* gene, which encodes a chaperone involved in mitochondrial protein homeostasis. Further analysis of a corresponding knock-in mouse model revealed that the *TRAP1* p.Q639* variant leads to male-specific behavioral abnormalities accompanied by altered neuronal plasticity and dendritic spine morphology typical for ASD.

## RESULTS

### A novel *TRAP1* p.Q639* variant in ASD-diagnosed monozygotic twin brother

We performed whole exome sequencing on three ASD-discordant male MZT pairs (Table S1) with DNA samples obtained from hair follicles. In ASD-affected twin, but not in his phenotypically normally developed brother, two potentially causal postzygotic mutations (also not observed in blood samples of twins’ parents; Fig. S1) were found: a nonsense mutation p.Q639* in *TRAP1* (hg19: chr16:g.003712013-G>A, NM_016292.2: c.1915C>T; VAF 8%) and *RUVBL1* p.F329L missense mutation (hg19: chr3:g.127816172-A>C, NM_003707.2: c.987T>G, p.F329L; VAF 48%) (Fig. 1A). Both TRAP1 and RUVBL1 variants have a frequency of 0 in the gnomAD database and our in-house database of over 10,000 Polish individuals who underwent exome sequencing.

**Figure 1.**
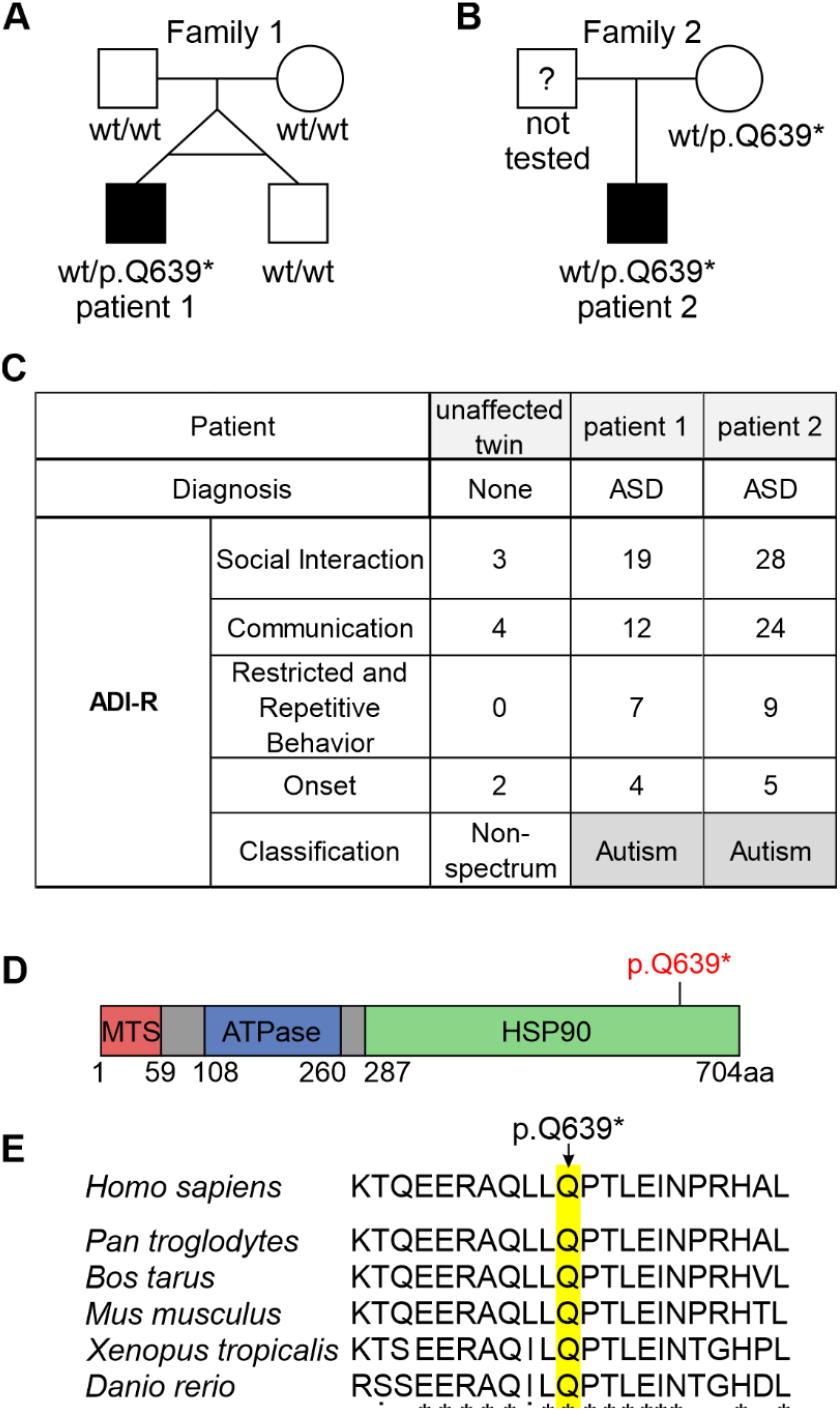
Identification of a *TRAP1* p.Q639* variant in ASD-affected individuals. Pedigree with phenotype and genotype denoted for (A) the examined MZT family and (B) the family of an additional ASD patient bearing the same *TRAP1* mutation from replication cohort; circle denotes female, square denotes male, symbol fill indicates ASD diagnosis. In a case of MZTs genotype derives from hair follicle DNA samples. (C) Results of the psychological evaluation of the ASD-unaffected twin, patient 1 (ASD-discordant MZT) and patient 2 (from replication cohort) using the Autism Diagnostic Interview-Revisited (ADI-R). (D) Diagram depicting the location of identified p.Q639* variant in the TRAP1 protein. (E) Comparative sequence alignment of the TRAP1 protein in different species shows universal Q639 residue conservation in these species (highlighted).

The *TRAP1* p.Q639* was *in-silico* predicted as pathogenic by meta scores predictors (including BayesDel addAF, BayesDel noAF, 8 points) and five individual predictors (including CADD phred, EIGEN, EIGEN PC, FATHMM-MKL, GERP RS), as variant of uncertain significance by three individual predictors (including DANN, LRT, MutationTaster), and as benign by FATHMM-XF. Moreover, the C nucleotide at position c.1915 is highly conserved (PhyloP100 9.434) and Q residue at position 639 is also located in an evolutionary conserved region of HSP90 domain of TRAP1 protein (Fig.1E). The *RUVBL1* p.F329L was predicted as benign by both meta scores and individual predictors. Furthermore, the T nucleotide at position c.987 is not conserved (PhyloP100 0.883). The p.Q639* variant introduces a premature stop codon, however the Z-score suggests that *TRAP1* gene is tolerant for loss-of-function (LoF) variants (−0.526). To assess the frequency of *TRAP1* LoF variants in the general population, we analyzed the distribution of LoF changes in the gnomAD database (v3.1.2). 61 different heterozygous LoF variants is reported in 202 individuals (total number of individuals > 150,000; ∼0.135%), including 42 (68,8%) located in the C-terminal HSP90 of TRAP1 protein (in 162 individuals, 80.2%). No significant difference in sex distribution was observed among *TRAP1* LoF variants carriers (95 males vs 107 females and 80 males vs 82 females when only HSP90 domain was considered).

Of note, in contrast to hair follicle DNA, both brothers were mosaic for the mutations in their blood samples (both with the same VAF: 2% for *TRAP1* variant and 22% for *RUVBL1*, respectively) (Fig. S1). Therefore, even though the *TRAP* and *RUVBL1* variants emerged post-twinning in the ASD-affected brother, they were detected in the phenotypically normal twin blood sample. This may reflect postulated blood chimerism in MZTs due to the transfer of hematopoietic stem cells between the twins in utero, which mask genetic differences between disease-discordant MZTs when mutational screening on blood-derived DNA (*18, 19*), underscoring the importance of using non-hematopoietic tissues (*20, 21*). Any genetic differences between MZTs are considered as postzygotic events. Since *TRAP1* p.Q639* mutation (VAF 8%) was found only in ASD-affected twin (when ectoderm-derived DNA was tested), thus it is considered as post-twinning event. The timing of mutation occurrence determines the variant distribution across different tissues (tissue specific mosaicism) and VAF (*22*). The role of postzygotic mutations in ASD is well established (*7-10, 23, 24*). For identification of causative mosaic mutations sequencing of DNA from affected tissue is recommended. The tissue which may be affected by postzygotic mutations in ASD patient is a brain available only post-mortem (*25, 26*). If affected tissue is inaccessible, then developmentally closest tissue that is available could be considered. Thus, in ASD-discordant MZTs we performed WES analysis using hair follicles derived from ectoderm, the same germ layer which contributes to brain. It was suggested that many postzygotic mutations with low VAF (>5%) observed in multiple tissues are likely to be mosaics in brain tissue as well (*27*).

To investigate the potential causative role of *TRAP1* and *RUVBL1* genes in influencing the risk of ASD, the entire coding sequence and intron-exon boundaries of both genes were analyzed in 176 unrelated ASD patients from Polish population (Table S2). No rare, functional variants were found in *RUVBL1*, while in four ASD patients variants in *TRAP1* were identified, including three heterozygous missense variants and one identical heterozygous nonsense p.Q639* *TRAP1* variant (VAF 50%). Interestingly, all *TRAP1* variants identified in the replication cohort of ASD patients were located within C-terminal HSP90 domain of TRAP1 protein (Figure 1D, Table S3).

In the case of the second male ASD patient which had a similar psychological evaluation as the first patient (Fig. 1C), the *TRAP1* p.Q639* variant was inherited from a heterozygous phenotypically normal mother (Fig. 1B). This could suggest sex-specific variable expressivity for this particular mutation. However, we could not establish the p.Q639* carrier status of the patient’s grandparents. Given the *TRAP1* truncation variant was found in more than one ASD patient, we focused on this gene.

Thus far, there is only one report linking *TRAP1* mutations with neurodevelopmental diseases, including ASD (*28*). Reuter at al., reported *TRAP1* homozygous splicing variant in patient with moderate intellectual disability, mental deterioration, autism, self-mutilation, muscular hypotonia, nystagmus and leukodystrophy. Loss-of-function variants in *TRAP1*, including nonsense variants, were described in one late-onset Parkinson’s disease patient (homozygous p.R47* (*29*), and in three patients with functional disorders (heterozygous p.E216*, p.Y229*, p.R703* (*30*). All previously reported nonsense variants are located within known functional domains: p.R47* is positioned within N-terminal mitochondria targeting sequence, p.E216* and p.Y229* within ATPase domain, and p.R703* within C-terminal HSP90 domain.

### Knock-in *Trap1* p.Q641* mice do not display a gross phenotype

The identified *TRAP1* p.Q639* mutation is predicted to generate a premature translation termination codon at position 639 of a mitochondrial chaperone of the HSP90 family. In order to investigate the potential mitochondrial etiology and sex-specificity of disease, we decided to employ an animal model of the *TRAP1* mutation. We generated a knock-in mouse with the identical *Trap1* mutation as that identified in two autistic patients (p.Q641* in mice is an equivalent of p.Q639* in humans). The mutation was introduced into the mouse genome using CRISPR-Cas9 (Fig. 2A,B). Both homozygous *Trap1*^*Q641*/Q641\**^ (MUT) and heterozygous *Trap1*^*WT/Q641\**^ (HET) animals were viable, fertile, and exhibited no morphological abnormalities, consistent with the phenotype of ASD patients and with a prior report that *Trap1* is not essential in mice (*31*). Moreover, the *Trap1* mutation did not affect overall brain morphology (Fig. S2). We next examined the expression of *Trap1* in mouse brains. Both mRNA and protein levels of *Trap1* were significantly reduced in heterozygotes, and truncated Trap1 protein was not detected in mutant mice (Fig. 2D,E).

**Figure 2.**
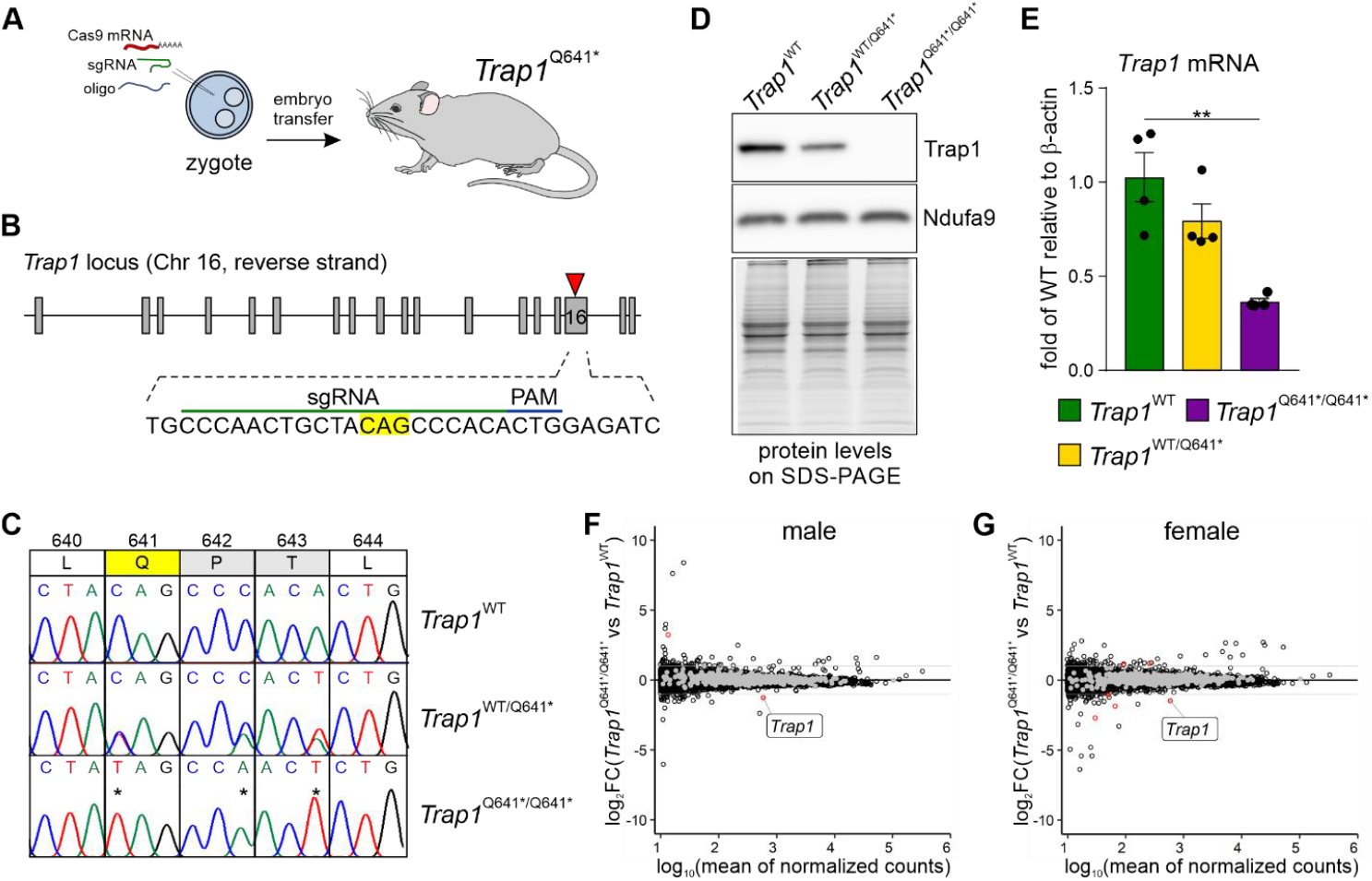
Generation and transcriptomic analysis of a mouse with the proband *Trap1* mutation (p.Q639*) equivalent. (A) Schematic of the strategy to introduce a knock-in mutation into the *Trap1* locus using CRISPR/Cas9. (B) Diagram of sgRNA targeting site in the mouse *Trap1* locus. Codon for Q641 is highlighted; PAM = protospacer adjacent motif. (C) Alignment of chromatograms covering the desired mutation site in *Trap1*: wild-type (*Trap1*^*WT*^, top), heterozygous (*Trap1*^*WT/Q641\**^, middle) and mutant (*Trap1*^*Q641*/Q641\**^, bottom). Mutated nucleotides are marked with asterisks. (D) Trap1 protein levels in the hippocampus of *Trap1*^*WT*^, *Trap1*^*WT/Q641\**,^ and *Trap1*^*Q641*/Q641\**^ mice. Ndufa9 as loading control and SDS-PAGE TGX-gel are shown. (E) *Trap1* mRNA levels in the cortex of *Trap1*^*WT*^, *Trap1*^*WT/Q641\**^ and *Trap1*^*Q641*/Q641\**^ mice as assessed by qPCR. Data is presented as fold change of *Trap1*^*WT*^, relative to *β-actin* mRNA; n=4 animals/group; **p<0.001, one-way ANOVA, *post-hoc* Dunnett’s multiple comparisons test; error bars indicate SEM. (F, G) MA plots representing the global differential gene expression in RNA-Seq analysis of the hippocampi of *Trap1*^*WT*^ and *Trap1*^*Q641*/Q641\**^ mice for male (F) and female (G) mice. The x-axis indicates the mean of normalized reads, and the y-axis indicates the log2 fold change of expression. Black circles represent transcripts not differentially expressed, red circles represent transcripts significantly differentially expressed (|log2(FC)| >1, p.adj-value < 0.05), and gray circles represent genes encoding proteins with mitochondrial localization. Differentially expressed transcripts in males: *Trap1* and *Xlr3a* and females: *Trap1, Gbp3, Miip, Rpl3-ps1, Gm15879, Gm49692*. n=3-4 animals/group.

TRAP1 has been suggested to be involved in the regulation of mitochondrial RNA processing (*32*). To obtain insight into how *Trap1* p.Q641* influences the cellular transcriptome, we performed sequencing of RNA isolated from hippocampi of WT, *Trap1*^WT/Q641*^ and *Trap1*^Q641*/ Q641*^ male and female mice. We observed no global effect of *Trap1* p.Q641* on the transcriptome, with only a few differentially expressed genes (DEGs) rising to the level of significance (Fig. 2F, G and Fig. S3C). The comparison of heterozygous *Trap1*^WT/Q641*^ to WT also showed only a few DEGs (Fig. S3A and B). Previously, inhibition of *Trap1* by a small molecule (TPP) was shown to induce the mitochondrial unfolded protein response (UPR) and mitochondrial pre-RNA processing defects in cancer cells (*32*), so we carefully analyzed mitochondrial RNAs within our dataset. As for the cytoplasmic mRNA, no dysregulation or evidence of mitochondrial stress on the transcriptome was detected (Fig. S3E-H). *Trap1* p.Q641* did not affect pre-RNA processing (Fig. S3D,I) or lead to the induction of transcription factors and kinases involved in the UPR (Fig. S3J,K). Together, our results showed that the *Trap1* p.Q641* mutation has a mild, if any, effect on mitochondrial RNA metabolism and does not cause obvious detrimental phenotypes in mice.

### Social interaction deficits in *TRAP1* p.Q641* males

To determine whether the mouse model recapitulates phenotypes relevant to ASD, we tested the mice for deficits in social interaction, one of the three ASD diagnostic criteria that can be replicated in mice (*33*). Both individuals carrying the *TRAP1* p.Q639* mutation and diagnosed with ASD showed abnormal social interactions in ADOS and ADIR tests. To assess the potential social impairment of the mice, we used Eco-HAB, an automated system for studying animal behavior (*34*)(Fig. 3A). The locomotor activity of *Trap1*^*WT*^, *Trap1*^*WT/Q641\**^, and *Trap1*^*Q641*/Q641\**^ mice in consecutive days was the same for all genotypes and was typical for mouse circadian rhythm (Fig. 3B,C). Analysis of in-cohort sociability, which measures how much time each pair of animals living in the same cage spends together, revealed deficits in social interaction in male homozygous and heterozygous mutant mice compared with WT (Fig. 3D). *Trap1* MUT and HET males tend to spend significantly less time in the group than WT males. No deficits in sociability were observed in *Trap1* mutant females. In fact, we observed slightly higher social interaction in *Trap1* HET mice as compared to WT (Fig. 3E), indicating that the *Trap1* p.Q641* mutation in mice results in social deficits, specifically in males. These results are consistent with the diagnosis of ASD in one patient from the replication cohort but not in his mother, despite the presence of the *TRAP1* p.Q639* mutation in both, suggesting that the effect of this mutation is sex-specific.

**Figure 3.**
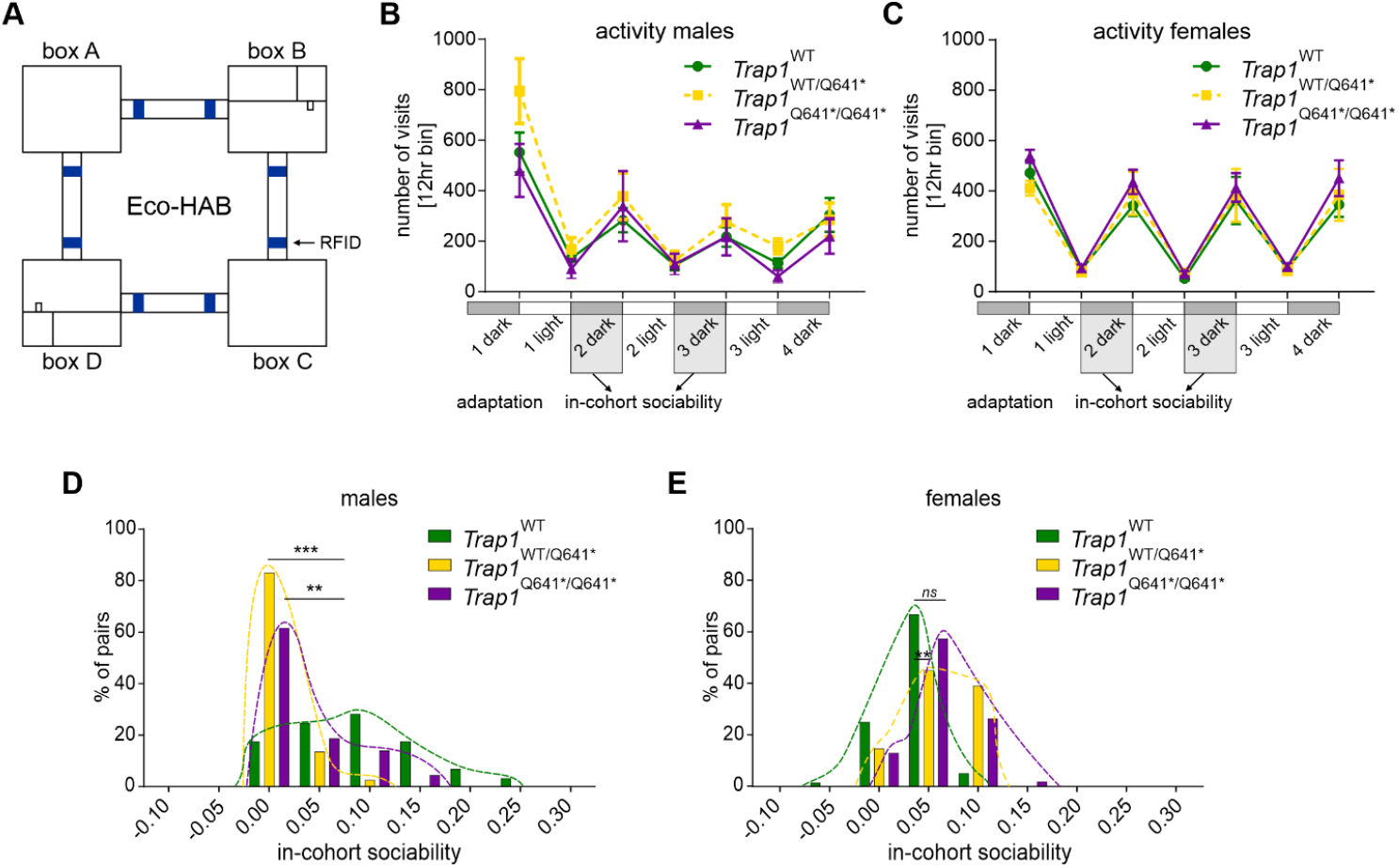
*Trap1* mutation in mice leads to deficits in social interaction in males. (A) Schematic of the Eco-HAB apparatus, which consists of four housing compartments linked together with tube-shaped passages, where RFID-antennas track mouse location. Food and water are available in two compartments: boxes B and D. (B, C) Analysis of the locomotor activity of *Trap1*^*WT*^, *Trap1*^*WT/Q641\**,^ and *Trap1*^*Q641*/Q641\**^ mice of both sexes. No significant differences were observed. (D, E) Histograms show the distribution of “in-cohort sociability” parameters for all pairs of animals in the *Trap1*^*WT*^, *Trap1*^*WT/Q641\**^, and *Trap1*^*Q641*/Q641\**^ cohorts in males (D) and females (E). In-cohort sociability measures how much time a pair of animals voluntarily spends together. Data is presented as a relative frequency distribution histogram; n=7-12 animals/group; ***p<0.001, **p<0.01; Kolmogorov-Smirnov test. See also Fig. S4.

### Increased spine density and basal synaptic neurotransmission in *Trap1*^*Q641*/ Q641\**^ males

The synaptic imbalance observed in ASD is thought to give rise to functional changes in neural circuitry caused by altered properties of individual synapses (*35*) located on dendritic spines undergoing plastic changes in response to synaptic stimulation. We analyzed spine morphology and electrophysiological recordings in the CA1 region of the hippocampus of WT, HET, and MUT male and female mice. For each mouse, one hemisphere was stained with DiI for assessment of the morphology of dendritic spines, while the second hemisphere was used for electrophysiological recordings. We observed striking differences between males and female brains in these assays. Male *Trap1*^*Q641*/Q641\**^ mice displayed increased spine density but spines were shorter and had decreased area (Fig. 4B-E). In contrast, the effect of the mutation was the opposite in females, with slightly reduced spine density and larger individual spines (Fig. 4F-J). The amygdala and medial prefrontal cortex (mPFC) are proposed to be one of several neural regions that are abnormal in autism (*36*). Therefore, we analyzed dendritic spine density and morphology in mPFC and amygdala of WT, HET and MUT male and female mice, where we also saw a sex-specific difference. No alterations were observed in male mPFC, except for increased spine density in heterozygous mice (Fig. S5A-E). In contrast, females showed increased spine density, length, head width, and area (Fig. S5F-J). In the amygdala, close to no differences between analyzed groups were observed (Fig. S5K-T).

**Figure 4.**
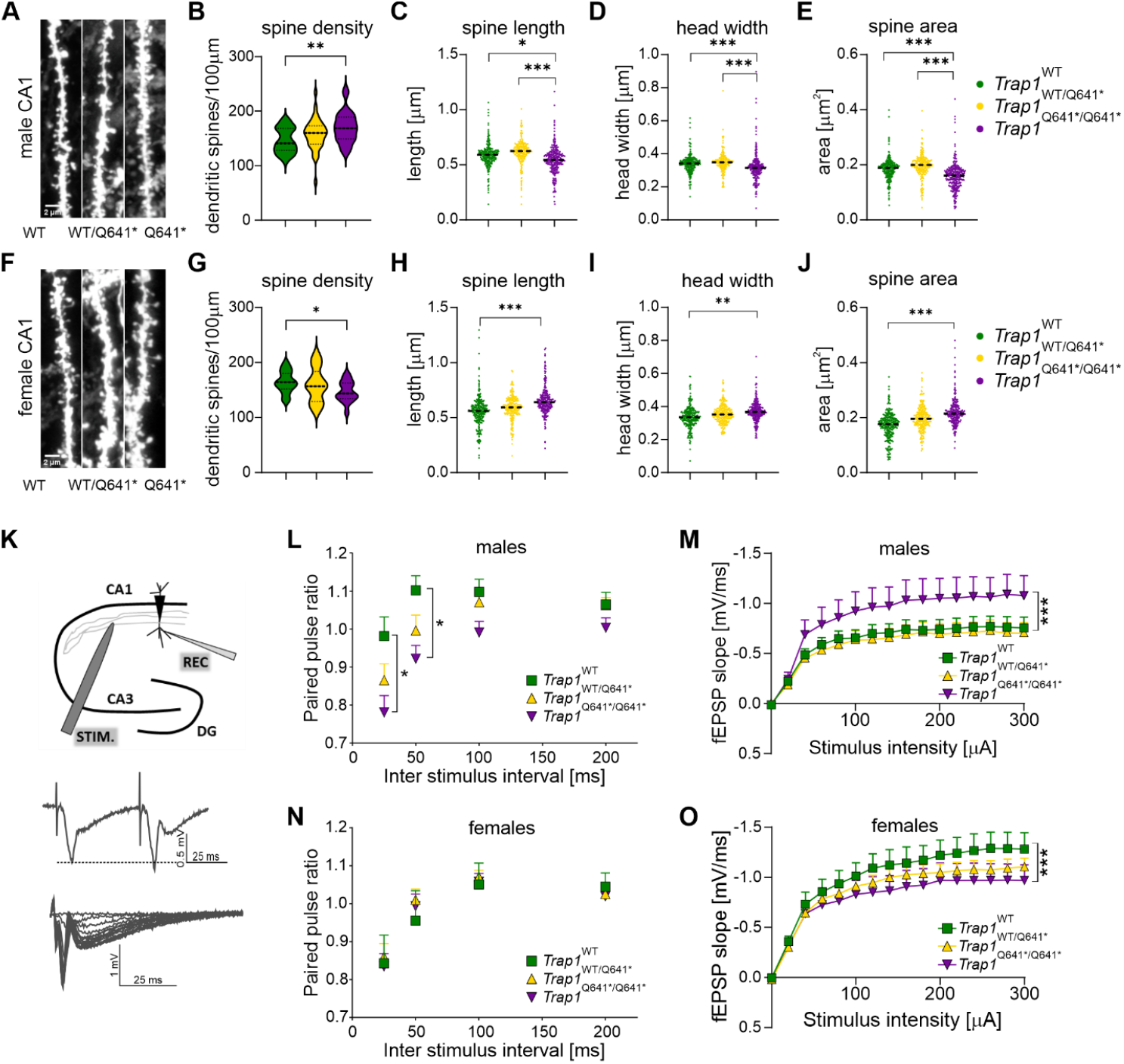
Altered dendritic spine density, morphology and excitatory synaptic transmission in *Trap1*^*Q641*/Q641\**^ mutant mice. (A, F) Representative images of DiI stained dendrites in the CA1 region of the hippocampus in males (A) and females (F). Scale bars 2µm. (B, G) The mean density of dendritic spines in the hippocampus in males (B) and females (G). Values show mean density/100 µm of dendrite; n=27-38 (males), n=21-31 (females) images/group; *p<0.05, **p<0.001; one way ANOVA, *post-hoc* Tukey’s test. (C-E, H-J) Dendritic spine morphology in *Trap1* male and female CA1 region of the hippocampus, including spine length (C, H), spine head width (D, I) and spine area (E, J); n=5274–8555 (males), n=4010-6118 (females) analyzed spines/experimental group; *p<0.05, **p<0.001, ***p<0.0001; nested ANOVA, *post-hoc* Tukey’s test. N=3-6 animals/group (males), N=3-4 animal/group (females). (K) Schematic of the electrophysiological recording setup depicting positions of stimulating (STIM) and recording (REC) electrodes in the CA1 hippocampal region. Middle, example trace of fEPSPs scaling in response to paired stimulation of Schaffer collaterals (inter stimulus interval 50ms). Bottom, example traces of compound fEPSPs were recorded in response to monotonically increasing stimuli applied to Schaeffer collaterals. (L) Male *Trap1*^*Q641*/****/****Q641\**^ mice exhibited significantly enhanced AMPAR-mediated synaptic responses recorded in response to monotonically increasing stimuli compared to *Trap1*^*WT/Q641\**^ and WT littermates (Kruskal-Wallis One Way ANOVA, p<0.001). (M) Female *Trap1*^*WT/Q641\**^ and WT littermates exhibited significantly enhanced AMPAR-mediated synaptic transmission compared to the *Trap1*^*Q641\*****/****Q641\**^ group but were not different from each other (Kruskal-Wallis One Way ANOVA, p<0.001, Dunn’s Method for multiple pairwise comparisons). (N) The average paired-pulse ratio for 25ms and 50ms inter-stimulus intervals was significantly reduced in the male *Trap1*^*Q641\*****/****Q641\**^ group compared to both *Trap1*^*WT/Q641\**^ and WT littermates (two-way RM ANOVA, p<0.01). (O) Average paired-pulse ratios were not significantly different in any of the female genotype groups (two-way RM ANOVA p>0.05). Males: N=3-6 animals, n=12-25 slices; Females: N=3-4 animals, n=12-17 slices. See also Fig. S6.

Using the same mice, we investigated the extent to which the TRAP1 mutation affects basal excitatory synaptic transmission and short-term forms of synaptic plasticity in the CA1 hippocampal region of mice. We hypothesized that the increased density of dendritic spines in male MUT mice could be reflected in changes of excitatory synaptic transmission. To test this possibility, we recorded AMPAR- and NMDAR-mediated field excitatory postsynaptic potentials (fEPSPs) in the CA1 hippocampal region. AMPAR-mediated synaptic signals were significantly more pronounced in the hippocampus of male homozygous *Trap1*^*Q641*/Q641\**^ mice compared with heterozygous and WT animals, consistent with the observed increased spine density (Fig. 4L). In contrast, female *Trap1*^*Q641*/Q641\**^ mice with lower dendritic spine density showed decreased synaptic transmission (Fig. 4M). The NMDAR-mediated component of fEPSP was most pronounced in male homozygous *Trap1*^*Q641*/Q641\**^ mice, whereas female *Trap1*^*Q641\**^ homo- and heterozygous mice exhibited a trend towards reduced synaptic transmission (Fig. S6 B,D). The composition of postsynaptic glutamate receptors plays an essential role in the plasticity of the local circuit. Altered availability of AMPARs and NMDARs in the synapse can lead to altered synaptic development, plasticity, and neuronal connectivity, resulting in synaptic pathophysiology (*37*). However, we found no changes in the sensitivity of fEPSPs to the AMPAR blocker DNQX in either sex of mouse regardless of genotype, suggesting that there is no gross change in the synaptic AMPAR/NMDAR ratio (Fig. S6 C,E).

Dysregulation of synaptic plasticity has been described in various animal models of autism (*38-40*). To examine short-term synaptic plasticity, we used the paired-pulse facilitation which results in scaling the synaptic responses proportional to initial probability of presynaptic release (*41*). We found that the average paired-pulse ratio for 25-ms and 50-ms interstimulus intervals was significantly reduced only in male *Trap1*^*Q641*/Q641\**^ mice but not in female mice (Fig.4 N,O).

Together, our electrophysiological and neuroanatomical data show that male *Trap1*^*Q641*/Q641\**^ mice specifically exhibit increased spine density and significant upregulation of basal synaptic neurotransmission in the hippocampus. Reduced paired-pulse ratios in these male MUT mice indicate decreased short-term synaptic facilitation. This may result from either increased basal neurotransmitter release probability (and thus reduced capacity for synaptic scaling) or impaired mechanism of the presynaptic release. In practice, reduced paired-pulse facilitation may decrease the ability of *Trap1*^*Q641*/Q641\**^ mice neurons to perform temporal summation of synaptic activity and propagate information following repetitive synaptic activity. Increase in the denstity of excitatory synapses, observed in *Trap1*^*Q641*/Q641\**^ male mice, could be a compensatory mechanism ameliorating this disturbance in presynaptic site.

### Fewer presynaptic mitochondria and changes in metabolic substrate usage in *Trap1*^*Q641*/Q641*\*^ males

The *Trap1*^*Q641*\****/****Q641\**^ mutation exhibits male-specific phenotypes in mice, resulting in a reduced ratio of paired impulses, an electrophysiological phenomenon characteristic of impaired presynaptic release of neurotransmitters. Since the proper functions of the presynaptic compartment rely on stable and functional mitochondria (*42*), we studied the CA1 region of the hippocampus in our mouse model using 3D electron microscopy. We examined the number, volume, and area of presynaptic mitochondria. Notably, we found that the density of presynaptic mitochondria was significantly lower in the CA1 hippocampus of male *Trap1*^*Q641*^ mice, both homozygous and heterozygous (Fig. 5A-G). However, the average size and volume of individual mitochondria remained the same (Fig. 5H, I). These results, along with our previous finding of higher dendritic spine density in these male mice (Fig. 4B), suggest that some synapses may lack mitochondria in the presynaptic compartment.

**Figure 5.**
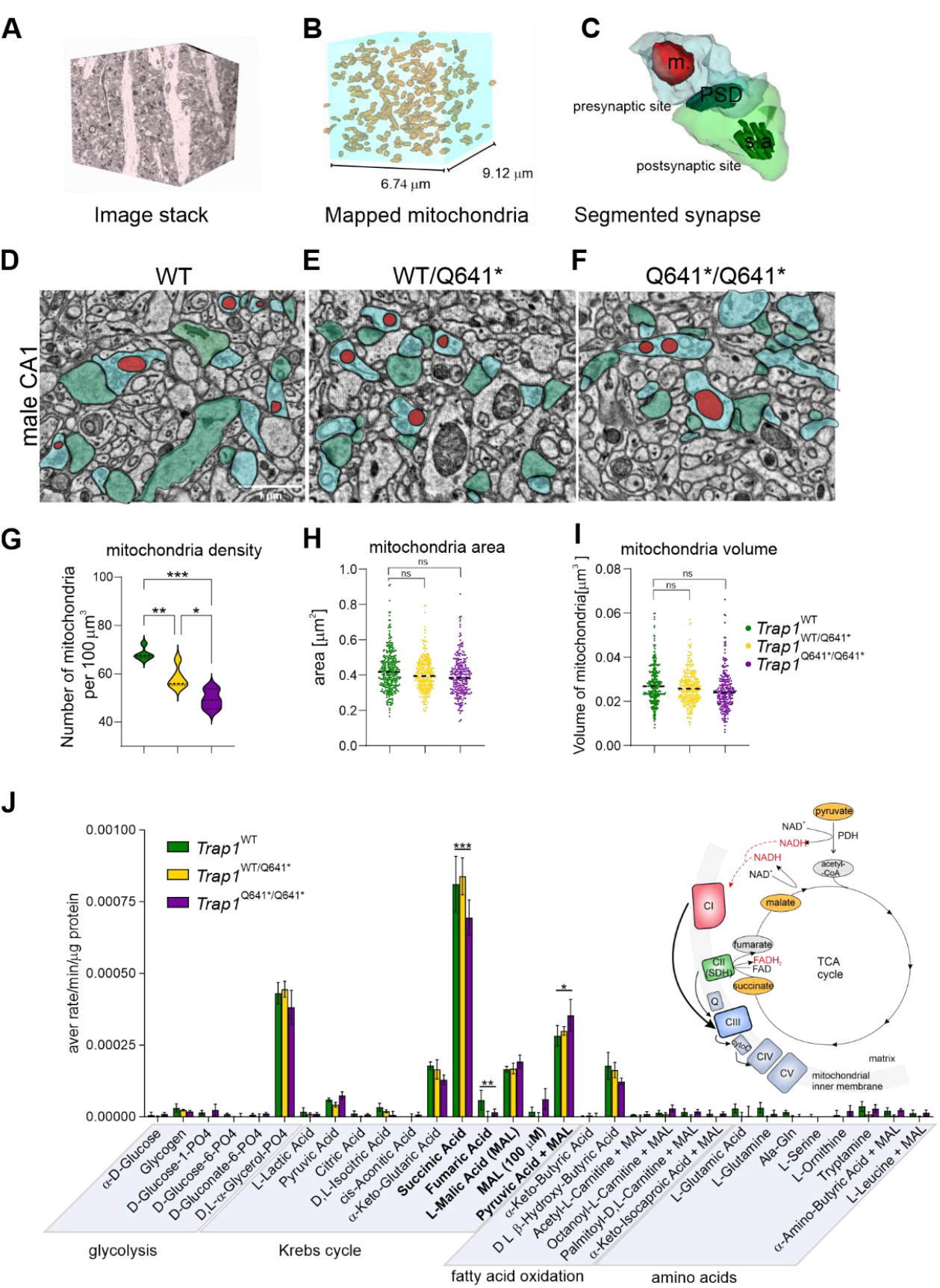
Decreased number of synaptic mitochondria in *Trap1*^*Q641\**^ and *Trap1*^*WT/Q641\**^ male mice. (A) Representative Serial Block Face Electron Microscopy (SBF-EM) image stack of stained striatum radiatum in the CA1 region of the hippocampus. (B) Illustration of mapped mitochondria in the image stack following manual segmentation. (C) An example of a segmented synapse with a presynaptic site (blue) with a mitochondrion (m, red), and a postsynaptic site (green) containing postsynaptic density (PSD) and spine apparatus (sa). (D-F) Representative electron micrographs of the CA1 region of the hippocampus in *Trap1* mice; pre- and postsynaptic sites with mitochondria are depicted as in C. (G) Decreased mitochondria density in *Trap1*^*Q641*/Q641\**^ and *Trap1*^*WT/Q641\**^ male mice. Plot shows mean density/100 µm^3^; N=3-5 animals/group; *p<0.05, **p<0.01, ***p<0.001; one way ANOVA, *post-hoc* Tukey’s. (H, I) No differences in mitochondria area (H) or volume (I) among tested genotypes. p>0.05, nested ANOVA. (J) Mitochondrial metabolism in synaptoneurosomes isolated from mouse brains of male *Trap1* mice. The rates of production of NAD(P)H and FADH_2_ from 31 different bioenergetic substrates, including glycolysis, TCA cycle intermediates, fatty acids and amino acids, were measured using MitoPlate™. Decreased utilization of succinate (***p < 0.001) was observed. Also, increased consumption of pyruvate + malate (*p < 0.05) was noticed. Results are presented as the average rate/min/μg of protein, +/-SEM (*n* = 3 per genotype; two-way ANOVA, *post hoc* Sidak’s multiple comparisons test). Scheme on the graph depicts selected substrate supply for mitochondrial respiration. Substrates differentially utilized in *Trap1*^*Q641*/Q641\**^ synaptoneurosomes are marked in *yellow*.

To further explore mitochondrial metabolism at synapses, we used a simple model system (synaptoneurosomes) and measured electron flow rates in the electron transport chain (ETC) from 31 different mitochondrial substrates.(Fig. 5J and Fig S7). Although there were no overall changes in mitochondrial metabolism (Fig. S7), we found significant differences in the use of the tricarboxylic acid cycle (TCA) substrates. Specifically, we observed decreased use of succinate, which produces FADH2 and donates electrons to the ETC at complex II in *Trap1*^*WT/Q641\**^ and *Trap1*^*Q641*/Q641\**^ males compared to WT littermates (Fig. 5J). At the same time, we detected increased consumption of pyruvate + malate, which produces NADH and donates electrons at complex I. Both complex I and complex II pass electrons to complex III. In this context, the reduced activity of complex II may be balanced by the increased activity of complex I to fuel the mitochondrial ETC.

## Conclusions

The link between ASD and mitochondria has been supported by clinical studies that reported disturbances at the levels of DNA, the activity of OXPHOS complexes, oxidative stress, and metabolites in the blood and urine of ASD patients (*43*). ASD and mitochondrial disorders share common clinical features, and the prevalence of mitochondrial disease is 500 times higher in the ASD population (*44*). However, only a handful of genes encoding proteins related to mitochondrial function have been directly shown to be associated with ASD. Three related genes – neurofilament light polypeptide (NEFL), mitochondrial uncoupling protein 4 (Slc25A27), and mitochondrial aspartate/glutamate carrier gene (Slc25A12) have significant genetic associations with ASD (*45-47*). Mitochondria are especially crucial in the context of the high energetic and mitochondrial demands of neurons. At the synapse, mitochondria are known to be essential for synaptic plasticity (*42*). In this study, we characterize a mutation leading to downregulation of expression of a mitochondrial chaperone protein, likely to cause mitochondrial impairment. This mutation, in the *TRAP1* gene is present in two unrelated male patients diagnosed with ASD, as well as in one unaffected mother. An interesting aspect of our study, using a newly generated mouse model harboring the identified *Trap1* p.Q639* variant, is the recapitulation of the sex-specific effects of the mutation. We detected not only male-specific behavioral abnormalities but also decreased number of presynaptic mitochondria in the male homozygous mutant brain, likely explaining their increased synapse strength and higher dendritic spine density. These synaptic aberrations in turn can explain deficits in social interaction exhibited by *Trap1* MUT male mice.

The etiology of ASD remains elusive, but recent studies, including the data presented, provide evidence for the role of mitochondrial homeostasis in autism. Thus, investigating strategies to modulate mitochondrial metabolism may hold promise for developing effective ASD treatments.

## References and Notes

## Acknowledgments

We thank prof. K. Radwanska, the Nencki Institute for granting access to the electrophysiology setup. NGS was performed thanks to Genomics Core Facility CeNT UW (RRID:SCR_022718), using NovaSeq 6000 platform financed by Polish Ministry of Science and Higher Education (decision no. 6817/IA/SP/2018 of 2018-04-10). Electron microscopy experiments were performed at the Laboratory of Imaging Tissue Structure and Function which serves as an imaging core facility at the Nencki Institute of Experimental Biology and is part of the infrastructure of the Polish Euro-BioImaging Node financed by Polish Ministry of Science and Higher Education contract no. 2022/WK/05; Polish Euro-BioImaging Node “Advanced Light Microscopy Node Poland”).

## Funding

This work was funded by the National Science Centre (NCN) Poland grant number 2014/13/B/NZ5/00287 to M.R., by NCN grant 2019/35/B/NZ4/04355 to M.D., TEAM TECH CORE FACILITY/2017-4/5 to AD and European Union’s Horizon 2020 research and innovation programme under grant agreement no 810425 to AD.

## Author contributions

Conceptualization: M.D., R.P., A.D.; Methodology: E.B., O.G., J.G.; Validation: All authors participated in the interpretation of the data; Formal analysis: M.R.,R.P, B.K., M.M., T.W., A.H., A.S., P.W., I.C.; Investigation: M.R., B.K., M.M., T.W., J.M., A.S., P.W., I.C.; Resources: R.P.; Writing - original draft: M.D., M.R., B.K.; Writing - review & editing: M.M., T.W., A.H., A.D., R.P.; Supervision: M.D., R.P., A.D., E.K.; Funding acquisition: M.R., M.D., A.D.

## Competing interests

Authors declare that they have no competing interests.

## Data and materials availability

All data are available in the main text or the supplementary materials. The mouse lines can be provided by A.D.’s pending scientific review and a completed material transfer agreement. Requests for the mice lines should be submitted to A.D. The whole exome sequencing data that support the findings of this study are available on request from the corresponding author R.P. The data are not publicly available due to ethical restriction (data contain information that could compromise the privacy of research participants). The RNA-Sequencing data discussed in this publication have been deposited in NCBI’s Gene Expression Omnibus and are accessible through GEO Series accession number GSE226319 (https://www.ncbi.nlm.nih.gov/geo/query/acc.cgi?acc=GSE226319).

## Notes

https://www.ncbi.nlm.nih.gov/geo/query/acc.cgi?acc=GSE226319

### Competing Interest Statement

The authors have declared no competing interest.

