## Supplementary material for "Mutation in mitochondrial chaperone TRAP1 results in male-specific autism"

**The PDF file includes:**

Materials and Methods
Figs. S1 to S6
Tables S1 to S3
References

#### Materials and Methods

Ethical statement

Material was collected from the ASD-discordant MZTs (blood and hair follicles), ASD affected patients (blood) and their parents (blood) after written informed parental consent was obtained. The study protocols were approved by the Institutional Review Board at Medical University of Warsaw (KB/153/2008 and KB/128/2014).

Clinical evaluation of monozygotic twins (MZTs) discordant for ASD

Phenotypically abnormal twins have received a clinical diagnosis of ASD determined by a multidisciplinary team including a psychiatrist based on ICD-10 diagnostic criteria. Clinical evaluation was performed through parent interviews and MZT examination using Autism Diagnostic Interview – Revised (ADI-R) (*1*, *2*) and Autism Diagnostic Observation Schedule (ADOS) (*3*) by a psychologist with experience in ASD diagnosis, as well as ADOS and ADI-R research reliability. All phenotypically abnormal twins met ADI-R and ADOS-2 criteria for an autism spectrum disorder.

DNA extraction and zygosity testing

DNA from whole blood was purified using the standard salt-out method. DNA from hair follicles was extracted using the DNA IQ™ Casework Pro Kit for Maxwell® 16 (Promega, Madison, WI, USA). Twin zygosity was determined using blood genomic DNA and analyzed with AmpFLSTR® NGM™ PCR Amplification Kit (Applied Biosystems, Foster City, USA) to assess 17 highly polymorphic markers. PCR products were separated on 31300xL Genetic Analyzer capillary sequencer (Applied Biosystems, Foster City, CA, USA) and evaluated using GeneMapper ID v3.2.1 (Applied Biosystems). Obtained results were analyzed according to allelic ladder standards.

Whole Exome Sequencing (WES)

WES analysis was performed using 50 ng of genomic DNA extracted from hair follicles using the SureSelectQXT Reagent Kit and SureSelectXT Human All Exon V5 (Agilent Technologies, Cedar Creek, TX, USA) according to manufacturer's instruction. Enriched libraries were paired-end sequenced (2x100 bp) on HiSeq1500 (Illumina, San Diego, CA, USA). For all samples > 50 million read pairs were generated resulting in average mean depth > 70x, coverage GE10 > 95% and GE20 > 90% of captured target.

WES sequencing data were analyzed as previously described (*4*). In brief, raw data was analyzed with bcl2fastq software (Illumina) to generate reads in fastq format. After the quality control step, including adapter trimming and low quality reads removal, reads were aligned to the GRCh37 (hg19) reference genome with Burrows-Wheeler Alignment Tool (http://bio-bwa.sourceforge.net/) and processed further by Picard (http://broadinstitute.github.io/picard/) and Genome Analysis Toolkit (https://software.broadinstitute.org/gatk/). Base quality score recalibration, indel realignment and duplicate removal were executed, and SNV and INDEL discovery was performed. Identified variants were further annotated with functional information, frequency in population (including EXaC, gnomAD, dbSNP, dbNSFP, 1000 genomes, as well as the frequency from in-house database of >10000 Polish individuals screened by WES), and known association with clinical phenotypes based on both ClinVar (*5*) and HGMD (*6*). Additionally, for enhancing the sensitivity of low allele fraction (possible mosaic state) GATK MuTect2 was used (*7*). Rare (frequency <0.01 in all tested databases) functional variants in protein-coding regions (missense, frameshift, stop-loss, stop-gain) and variants in splicing regions were considered. Priority was given to variants presented only in phenotypically abnormal twin (while in phenotypically normal co-twin lack of the variant was confirmed if covered >20x) or variants shared between twins but with significantly different variant allele fraction (effect of mutation load, incomplete penetrance). *In silico* pathogenicity prediction was performed based on Varsome (https://varsome.com) provided pathogenicity and conservation scores. All prioritized variants were manually inspected in Integrative Genomics Viewer (*8*).

Validation of single nucleotide variants (SNVs)

Detected SNVs observed in the ASD diagnosed twin but not in ASD-unaffected twin sibling were further validated by NGS-based deep amplicon sequencing (DAS). For validation analysis DNA samples isolated from blood (parents and MTZs) and hair follicles (MTZs) were tested. PCR primers for DAS were designed using Primer3 (http://primer3.ut.ee/) and are available upon request. Additionally, for DAS to the *locus*-specific primers overhang adapters were added (forward overhang: 5'-TCGTCGGCAGCGTCAGATGTGTATAAGAGACAG, reverse overhang: 5'-GTCTCGTGGGCTCGGAGATGTGTATAAGAGACAG) to make the PCR product compatible with the Nextera XT index primers (Illumina). Amplicons targeted identified SNVs were paired-end sequenced (2x100 bp) on an Illumina HiSeq1500.

Sequencing of *RUVBL1* and *TAP1* genes in a cohort of Polish ASD patients

For the replication study, the entire coding sequence and intron-exon boundaries of *RUVBL1* and *TRAP1* genes were analyzed using the NGS-DAS strategy in 176 unrelated ASD patients from the Polish population. Primers and PCR conditions are available upon request. For each patient, after PCR amplicon pooling, samples were subjected to NGS library preparation protocol using Nextera XT DNA Library Preparation Kit (Illumina), paired-end sequenced (2x250 bp) on Illumina MiSeq and analyzed as described above for WES. Additionally, from the in-house WES database (> 10000 samples) 100 unrelated, healthy adult men from Polish population were selected as a control group, and rare SNVs distribution in *RUVBL1* and *TRAP1* genes were analyzed.

Designing mice with point mutation in *Trap1* gene (*Trap1*(p.Q641*))

A new mouse line B6.CBA $Trap1^{\mathrm{em1Iimcb}}$/Tar ($Trap1^{Q641*}$) was generated by the Mouse Genome Engineering Facility (www.crisprmice.eu). All mice were bred and maintained in the animal house of

Faculty of Biology, University of Warsaw under a 12-h light/dark cycle with food and water available ad libitum. The animals were treated in accordance with the EU Directive 2010/63/EU for animal experiments.

Based on the mouse genome (GRCm38/mm10 Assembly) a single guide RNA (sgRNA) was designed using an online CRISPR tool (http://crispr.mit.edu). The chosen sequence did not show any major off-targets but had high calculated efficiency. For sgRNA synthesis two oligodeoxynucleotides (ODN) carrying T7 polymerase promoter, guide sequence (TRAP1_sgRNA) and sgRNA scaffold (Univ_IVsgRNA_2R) were used to form dsDNA template for *in vitro* transcription using in-house T7 Polymerase. A 120 bp oligonucleotide Trap1_mut_O1 carrying point mutations to replace the wild-type Trap1 fragment was designed to introduce c.1921C>T mutation (p.Q641*) at the end of exon 16 of *Trap1*. Additionally two silent mutations: c.1926C>A (p.P642P) and c.1929A>T (p.T643T) were introduced, to minimize the risk of cleavage of recombinant DNA by Cas9. Cas9 mRNA was in-vitro transcribed from Addgene pX458 plasmid using T7 RNA Polymerase. Poly(A) tail was added using E.coli PolyA Polymerase (Thermo Scientific) and m7Gppp5´N Cap was added with Vaccinia Capping System (NEB M2080) according to the manufacturer's protocols. Injection cocktail: (12.5ng/ul Trap1_sgRNA IVT, 25ng/ul Cas9 mRNA IVT, 1.5 pM Trap1_mut_O1) was introduced into mice zygotes via microinjections. After 24-48h of incubation embryos were implanted into surrogate mice.

Genotyping

Pups were genotyped at 4 weeks of age. DNA from tail or ear tips was isolated with Genomic Mini kit (A&A Biotechnology). gDNA was amplified with Trap1_Seq1F/1R primer pair using Phusion HSII polymerase and HF buffer, then the amplicons were sequenced.

PCR cycling:

98’C – 3:13

35x 98’C – 13s

63’C - 17s

72’C – 7s

72’C – 5:00

12’C - forever

Primers used in the study:

sgRNA sequence: 5’CCAACTGCTACAGCCCACAC3’

Trap1_Seq1F: 5’TGTGTACCCTTGGACATCTT3’

Trap1_Seq1R: 5’GAGAATGTCAGGTTTGTGCT3’

Trap1_mut_O1 (lowercase point mutations indicated): 5’ggagaccctgacaggtgtgtgctggagcccaggcctacCTGGGGTTGATCTCCAGaGTtGGCTaTAGCAGTTGGGCACGTTCCTCCTGGGTCTTGGCCAGCTGCTGCATACGCAAGAAAT3’

Trap1_sgRNA1F (gRNA sequence in red):
5’gaaatTAATACGACTCACTATAGGGCCAACTGCTACAGCCCACACGTTTTAGAGCTAGAAATAGCAAGTTAAAATAAGGC 3’
Univ_IVsgRNA_2R
5’AAAAGCACCGACTCGGTGCCACTTTTTCAAGTTGATAACGGACTAgccttattttaacttgctatttctagctcta 3’

Animals

Mice were bred in the Animal House of the Faculty of Biology, University of Warsaw. The animals were kept in the laboratory animal facility under a 12-h light/dark cycle with food and water available ad libitum. The animals were treated in accordance with the EU Directive 2010/63/EU for animal experiments and Polish regulations. All experimental procedures were pre-approved by the Local Ethics Committee (WAW/194/2016 and WAW/771/2018).

Tissue preparation

Young adult mice (~3 months old) were sacrificed and the hippocampi and cortices were dissected and frozen at -80°C for further investigation.

SDS-PAGE and western blotting

Hippocampi were homogenized in RIPA buffer using a Dounce homogenizer. Protein content was measured using Pierce BCA protein assay kit. Equal amounts of samples were resolved by SDS-PAGE (10% or 12% TGX Stain-Free FastCast Acrylamide Solutions, BioRad). After electrophoresis, proteins in the gel were visualized using Bio-Rad’s ImageLab software to verify equal protein loading. Proteins were transferred to PVDF membranes (pore size 0.45 μm, Immobilon-P, Merck Millipore) using Trans-Blot Turbo Blotting System (BioRad; 170-4155). Membranes were blocked for 1 h at room temperature in 5% non-fat dry milk in PBS-T (PBS with 0.01% Tween-20), followed by overnight incubation at 4°C with primary antibodies in 5% milk in PBS-T (anti-Trap1, BD Biosciences Cat# 612344, RRID:AB_399710; anti-Ndufa9, Abcam Cat# ab14713, RRID:AB_301431). Blots were washed 3 × 5 min with PBS-T, incubated 1 h at room temperature with HRP-conjugated secondary antibody (1:10,000 in 5% milk) and washed 3 × 5 min with PBS-T. HRP signal was detected using Amersham ECL Prime Western Blotting Detection Reagent (GE Healthcare) on Amersham Imager 600 using automatic detection settings.

RNA isolation and qRT-PCR

RNA was extracted from the mouse cortex using TRIzol (Thermo Fisher Scientific), DNA contamination was removed by 2 U of TURBO DNase (AM2238, Ambion) in the supplied buffer in 37°C for 30 min. Next, RNA was re-isolated with phenol/chloroform, precipitated with ethanol, and resuspended in 50 μl of RNase free water. RNA concentration was calculated from absorbance at 260 nm using a DS-11 Spectrophotometer (DeNovix). Next, equal amounts of RNA samples (1.1 μg) were reverse transcribed using random primers (GeneON; #S300; 200 ng/RT reaction) and SuperScript IV Reverse Transcriptase (Thermo Fisher Scientific). Subsequently, the cDNA samples were amplified using sequence-specific primers in a final reaction volume of 15 µl, using PowerUp SybrGreen MasterMix in a LightCycler480 (Roche). Two different primer pairs were used to analyse *Trap1* mRNA: Trap1-1 Fw: TCCGCAGCATCTTCTATGTG; Trap1-1 Rev: TATACAGTGCCACGCTGGAG; Trap1-2 Fw: CAGGACAGTTATACAGCACACAG; Trap1-2 Rev: CTCATGTTTGGAGACAGAACCC. Fold changes in expression were determined using the ΔΔ Ct (where Ct is the threshold cycle) relative quantification method. Values were normalized to the relative amounts of *Actb* mRNA. Primer sequences: Actb Fw: CCCAGAGCAAGAGAGGTATC; Actb Rev: ATTGTAGAAGGTGTGGTGCC.

RNA isolation, library preparation and RNA-sequencing

Total RNA was extracted from the mouse hippocampi with TRI Reagent (Sigma-Aldrich, Cat# 93289) according to the manufacturer's instructions and followed by DNase treatment (Invitrogen, Cat# 18080085). RNA quality and integrity was verified using RNA Pico 6000 (Agilent, Cat# 5067-1513). Strand-specific RNA libraries were prepared using a TruSeq Stranded Total RNA Library Prep (Illumina, Cat# 20020596) and adapters IDT for Illumina TruSeq RNA UD Indexes (96 Indexes, 96 Samples) (Illumina, Cat# 20022371) according to manufacturer's instructions. For library preparation 1µg of total RNA was used, fragmented by 8 minutes incubation at 94°C. The library was enriched with 11 amplification cycles. The quality of the enriched library was verified using 2100 Bioanalyzer and High Sensitivity DNA Kit (Agilent, Cat# 5067-4626). The libraries' concentration was estimated by qPCR means with KAPA Library Quantification Kit (Kapa Biosciences, Cat# KK4824), according to manufacturer's instructions. These libraries were subsequently sequenced using an Illumina NovaSeq 6000 sequencing platform and NovaSeq 6000 S1 Reagent Kit (200 cycles) (Illumina, Cat# 20012864) in 2x100nt pair-end mode with standard procedure according to manufacturer's instructions.

Quality control and mapping of highthrougput RNA sequencing data

The Illumina sequencing reads were quality filtered using Cutadapt v1.18 (https://www.doi.org/10.14806/ej.17.1.200 https://cutadapt.readthedocs.io/en/stable/ ) to remove Illumina adapter sequences, trim low-quality fragments (minimum Q score = 20), and remove reads that were shorter than 30 nt after trimming. After trimming, reads were repaired with bbmap v38.86 (https://doi.org/10.1371/journal.pone.0185056 BBMap – Bushnell B. – sourceforge.net/projects/bbmap/ ). Repaired reads were mapped to mouse genome (GRCm38) using STAR v2.7.5c (https://doi.org/10.1002/0471250953.bi1114s51 https://github.com/alexdobin/STAR ). For downstream analysis, only uniquely mapped reads were used. The feature assignments of reads was performed with HTSeq v0.12.4 (https://doi.org/10.1093/bioinformatics/btac166 https://github.com/htseq/htseq), choosing genes as representative features. The normalization and differential gene expression analysis was performed with DESeq2 R package (doi: 10.1186/s13059-014-0550-8 ). For mitochondrial junction analysis a custom annotation of junctions was generated based on mitochondrial gene annotation from Gencode basic annotation v25. The count of junctions were generated as for gene features with HTSeq. For RNA-Seq analysis all plots were generated in R/Bioconductor environment with the usage of ggplot2 package (Wickham H (2016). *ggplot2: Elegant Graphics for Data Analysis*. Springer-Verlag New York. ISBN 978-3-319-24277-4, https://ggplot2.tidyverse.org. ). The RNA-Sequencing data discussed in this publication have been deposited in NCBI's Gene Expression Omnibus and are accessible through GEO Series accession number GSE226319.

<u>Nissl-staining</u>

Mouse brains (from 3 month old mice) were fixed in 4% paraformaldehyde in PBS overnight at 4°C, then the brains were cryoprotected in 20% sucrose in PBS at 4°C for 48 hours and frozen in -80°C. Next, the brains were cut coronally on cryostat (Cryostat Leica CM 1860) on 40-μm slices. Coronal sections were air dried on slides and stained with 0.1 % cresyl violet solution (containing 3 % acetic acid) for 5 min, washed, dehydrated, cleared in xylene, and coverslipped.

<u>Eco-HAB</u>

Eco-HAB experiments were performed on young-adult male and female mice (~2.5- to 4-month-old). Experiments were performed as previously described (*9*). To individually identify animals in Eco-HAB, all mice were subcutaneously injected with glass-covered microtransponders (9.5 mm length, 2.2 mm diameter, RF*IP* Ltd) under brief isoflurane anaesthesia. Microtransponders emit a unique animal identification code when in range of RFID antennas. After injection of transponders, subjects were moved from the housing facilities to the experimental rooms and adapted to the shifted light/dark cycle of their new environment (the dark phase shifted from 20:00 – 8:00 to 13:00 – 01:00 or 12:00 – 24:00 depending on summer/winter UTC+01:00). For 2 weeks prior to behavioural testing, subjects were housed together and grouped appropriately for their respective experiment. The following cohorts were used: (a) males: WT, n=8; HET, n=9; MUT, n=7, (b) females: WT, n=11; HET, n=12; MUT, n=10. Cohorts were subjected to 84-hour Eco-HAB testing protocols (see Figure 3A and Figure S4 A) divided into an adaptation phase (24 h) and in-cohort sociability testing phase (next 48 h). Throughout the experiment mice could freely explore all compartments, with unrestricted access to food and water in two of the housing compartments. All the details concerning Eco-HAB apparatus construction, software package for data collection, processing and analysis is described in the original paper (*9*). Briefly, activity was defined as the number of visits of an animal to all of Eco-HAB compartments in 12 h bins. In-cohort sociability of each pair of mice within a given cohort is a measure of sociability that is unique to the Eco-HAB system. For each pair of subjects within a cohort, the times spent by the mice in each of the four compartments and the total time spent by the pair together in each of the cages were calculated. The in-cohort sociability was then defined by the total time spent together minus the time animals would spend together assuming independent exploration of the apparatus.

DiI staining of brain slices

To visualize changes in the shape of dendritic spines, 1,1′-dioctadecyl- 3,3,3,3′-tetramethylindocarbocyanine perchlorate (DiI) staining was performed in brain sections from *Trap1* mutant mice, heterozygotes and wild-type mice. The mice were anesthetized and transcardially perfused with 1.5% paraformaldehyde. The brains were dissected and sliced using a vibratome. Slices (100 µm thick) that contained the hippocampus were allowed to recover for at least 1 h at room temperature. Random dendrite labeling was performed using 1.6 µm tungsten particles (Bio-Rad, Hercules, CA, USA) that were coated with propelled lipophilic fluorescent dye (DiI; Invitrogen) that was delivered to the cells by gene gun (Bio-Rad) bombardment. Images of dendrites in hippocampal CA1, medial prefrontal cortex and amygdala were acquired under 561 nm fluorescent illumination using a confocal microscope Axio Imager Z2 Zeiss LSM 700 (63x objective, 1.4 NA) at a pixel resolution of 1024 x 1024 with a 3.4 zoom, resulting in a 0.07 µm pixel size. Images were acquired blinded to the genotype group.

Morphometric Analysis of Dendritic Spines

The analysis of dendritic spine morphology and calculation of changes in spine shape parameters were performed as described previously (*10, 11*). The images acquired from the brain slices were processed using ImageJ software (National Institutes of Health, Bethesda, MD, USA) and analyzed semi-automatically using custom-written SpineMagick software (patent no. WO/2013/021001) blinded to the genotype group. The analyzed dendritic spines belonged to secondary and ternary dendrites. We used length, head-width, area and a scale-free parameter of relative changes in the spine length-to-head width ratio, which reflects spine shape. The spine length was determined by measuring the curvilinear length along a fitted virtual skeleton of the spine. The fitting procedure was performed by looking for a curve along which integrated fluorescence was at a maximum. The head width was defined as the diameter of the largest spine section while excluding the bottom part of the spine (1/3 of the spine length adjacent to the dendrite). Dendritic segments of at least 3 animals per group were morphologically analyzed resulting in 4010 – 8555 spines. To determine spine density, approximately 1500 µm of dendritic length was analyzed per experimental group. For the statistical analysis of synaptic plasticity (density and morphology), we used the Nested-one-way ANOVA and Tukey's multiple comparisons test. Values of $p < 0.05$ were considered statistically significant. The analyses were performed using Prism 9.3.1 software (GraphPad, San Diego, CA,USA).

Electrophysiology recordings

Acute brain slices were prepared as described previously (*12*). All of the experimental procedures were approved by the Local Ethics Committee. The animals were anesthetized with isoflurane and decapitated. The hippocampi were dissected and cut into 350 µm thick slices using a vibratome (VT1200S, Leica, Germany) in ice-cold buffer that contained 75 mM sucrose, 87 mM NaCl, 2.5 mM KCl, 1.25 mM $NaH_2PO_4$, 25 mM $NaHCO_3$ , 0.5 mM $CaCl_2$, 10 mM $MgSO_4$*$7H_2O$, and 20 mM glucose, pH 7.4. Slices were subsequently allowed to recover in the same solution (32°C, 15 min) and stored until the end of the experiments in the oxygenated artificial cerebrospinal fluid (aCSF) that contained 125 mM NaCl, 25 mM $NaHCO_3$, 2.6 mM KCl, 1.25 mM $NaH_2PO_4$, 2.0 mM $CaCl_2$, and 20 mM glucose, pH 7.4. All solutions were oxygenated with carbogen (95% $O_2$ , 5% $CO_2$). Recordings were made in aCSF after 2 hours of slice recovery. Schaeffer collateral axons were stimulated with a concentric bipolar electrode (0.1 Hz, 0.3 ms). Compound AMPAR- and NMDAR-mediated fEPSPs were recorded with glass micropipettes that were filled with aCSF (1-3 M Ω resistance) in the stratum radiatum of the CA1 region (150 – 200 µ m from the stratum pyramidale). NMDAR-mediated signals were isolated from compound fEPSPs with the AMPA/kainate receptor antagonist DNQX (20 µM) and L-type calcium channel blocker nifedipine (20 µM) in $Mg^{2+}$-free solutions, as described previously (*12*). At the end of each recording, the NMDAR antagonist APV (50 µM) was used to confirm the origin of the recorded fEPSPs. All of the drugs were obtained from Sigma-Aldrich (Poland) and Tocris (UK). The electrophysiology data were analyzed using pClamp10.7.0.3 software (Molecular Devices, USA) and AxoGraphX software (developed by John Clements) as described previously (*12*). Recordings were performed blinded to the genotype group.

Serial Block-Face Scanning Electron Microscopy (SBEM)

The mice were transcardially perfused with 2% PFA (P6148 Sigma-Aldrich) with 2% glutaraldehyde (GA, EM grade, G5882 Sigma-Aldrich) in 0.1 M phosphate buffer pH 7.4 in ddH2O. The brains were gently dissected from the skull and fixed for SBEM overnight at 4 °C. Next, the brains were cut into 100 µm slices and the hippocampi were dissected for the staining (blade travel speed: 0.075 mm/s, cutting frequency: 80 Hz). The SBEM staining was performed according to previously published protocol (Corpus ID: 136219796). Slices were postfixed with solution of 2% osmium tetroxide (#75632 Sigma-Aldrich) and 1.5% potassium ferrocyanide (P3289 Sigma-Aldrich) in 0.1 M phosphate buffer pH 7.4 for 60 min on ice. Next, samples were rinsed 5 × 3 min with ddH2O and subsequently exposed to 1% aqueous thiocarbohydrazide TCH (#88535 Sigma) solution for 20 min. Samples were then washed 5 × 3 min with ddH2O and stained with osmium tetroxide (1% Osmium Tetroxide in ddH2O) for 30 min RT. Afterwards, slices were rinsed 5 × 3 min with ddH2O and incubated in 1% aqueous solution of uranyl acetate overnight in 4 °C. The next day, lead aspartate solution was prepared by dissolving lead nitrate (0.066 g) in 10 ml L-aspartic acid (0.998 g of L-aspartic acid (Sigma-Aldrich) in 250 ml of ddH2O). Slices were rinsed 5 × 3 min with ddH2O, incubated with lead aspartate for 30 min in 60 °C and then washed 5 × 3 min with degassed (autoclaved) ddH2O and dehydration was performed using graded dilutions of ethanol (ice-cold for better membrane preservation, 30%, 50%, 70%, 80%, 90%, and 2 × 100% ethanol, 5 min each). Samples were infiltrated with a resin that was prepared by mixing: A (17 g), B (17 g) and D (0.51 g) components of Durcupan (#44610 Sigma-Aldrich) with 8 drops of DMP-30 (#45348 Sigma) accelerator (*13*). Part of resin was then mixed 1:1 (v/v) with 100% ethanol and slices were incubated in 50% resin for 30 min in RT. The resin was then replaced with 100% Durcupan for 1 h in RT and then 100% Durcupan infiltration was performed o/n. The next day, samples were infiltrated with freshly prepared resin (as described above) for another 2 h in RT, then flat embedded between Aclar sheets (Ted Pella #10501-10). Samples were put in a laboratory oven for at least 48 h, 65–70 °C- for the resin to polymerize. After resin hardening, Aclar layers were separated and the resin embedded samples were taken out. Squares, of approximately 1 mm × 1 mm, cut out with razor-blades, were attached to aluminum pins (Gatan metal rivets, Oxford instruments) with a very small amount of cyanacrylate glue and then mounted to the ultramicrotome (Leica ultracut R) and trimmed. Samples were grounded with conductive silver paint (Ted Pella, 16062-15) to the pin and mounted into the 3 View chamber.

<u>3 View imaging, scan processing, image analysis and statistical analysis of the data</u>

Samples were imaged with SigmaVP (Zeiss) scanning electron microscope equipped with 3 View 2 chamber using a backscatter electron detector. Scans were taken in the middle part of stratum radiatum of the CA1 of the dorsal hippocampus. From each sample 200 sections were collected (thickness 60nm). Imaging settings: variable pressure18Pa, EHT 4 kV, aperture: 15 µm, pixel dwell time: 7 µs, pixel size: 5 nm (2048 × 2048 resolution). Obtained scans were aligned using the ImageJ software (ImageJ-> Plugins-> Registration-> StackReg) and saved as.tiff image sequence. Then the image sequences were imported to the Reconstruct software (*14*) (http://synapses.clm.utexas.edu/tools/reconstruct/reconstruct.stm). Mitochondria density was analysed with the modified unbiased brick method (*15*) per tissue volume. For each sample all mitochondria were counted in 1 brick. Size of each brick was 6.74 µm × 6.74 µm × 9.12 µm. A structure was considered to be a synaptic mitochondria when it was in a presynaptic site containing vesicles opposing postsynaptic site with electron dense material. Image acquisition and analysis was performed blinded to the genotype group. Statistical tests were performed in Graphpad Prism 9.5.0. Details of statistical tests are provided in figure legends. Mitochondria volume and area followed normal distributions and were compared with Nested one-way ANOVA test. Mitochondria density followed normal distribution and was compare with ordinary one-way ANOVA, pairs of results were compared with Tukey's multiple comparisons test. Differences between groups were considered significant if $p < 0.05$. Mean values and Standard Errors of the Mean (SEM) are shown.

<u>Preparation of synaptoneurosomes and MitoPlate™ Assay</u>

Synaptoneurosomes were prepared as described previously (*16*). Before tissue dissection Krebs buffer (2.5 mM $CaCl_2$, 1.18 mM $KH_2PO_4$, 118.5 mM NaCl, 24.9 mM $NaHCO_3$, 1.18 mM $MgSO_4$, 3.8 mM $MgCl_2$, 212.7 mM glucose) was aerated with an aquarium pump for 30 min at 4°C. Next, the pH was lowered to 7.4 using dry ice. The buffer was supplemented with 1×protease inhibitor cocktail cOmplete EDTA-free (Roche) and 60 U/ml RNase Inhibitor (RiboLock, Thermo Fisher Scientific). Animals were euthanized by cervical dislocation, hippocampi and a part of cortex adjacent to the hippocampus were dissected. Tissue from one hemisphere (~50 mg) was homogenized in 1.5 ml Krebs buffer using a Dounce homogenizer with 10-12 strokes. All steps were kept ice-cold to prevent stimulation of synaptoneurosomes. Homogenates were loaded into 20 ml syringe and passed through a series of pre-soaked (with Krebs buffer) nylon mesh filters consecutively 100, 60, 30 and 10 µm (Merck Millipore) in cold room to 50 ml polypropylene tube, centrifuged at 1000 g for 15 min at 4°C, washed and pellet was resuspended in Krebs buffer with protease and RNase inhibitors. Mitochondrial function assays using MitoPlates S-1 (Biolog, Cat. #14105) were performed as suggested by the manufacturer. Briefly, Assay Mix was dispensed into all wells of a MitoPlate and the plate was incubated at 37°C for 1 hour to allow substrates to fully dissolve. Freshly isolated synaptoneurosomes were pelleted and resuspended in 1xBiolog Mitochondrial Assay Solution (MAS). SN suspension was dispensed into each well of a MitoPlate (30 µl/well) and the color formation at 590 nm was read kinetically for 2 hours on a Multiskan FC Microplate Photometer (Thermo Scientific). The background was corrected for the blank sample and the average rate between 10 and 100 min was calculated. The protein content of synaptoneurosomal samples was measured using Bradford method. Synaptoneurosomes isolated from 3 male and 3 female mice per genotype were analyzed. Results are presented as average rate per minute per µg of protein.

#### Supplementary Figures

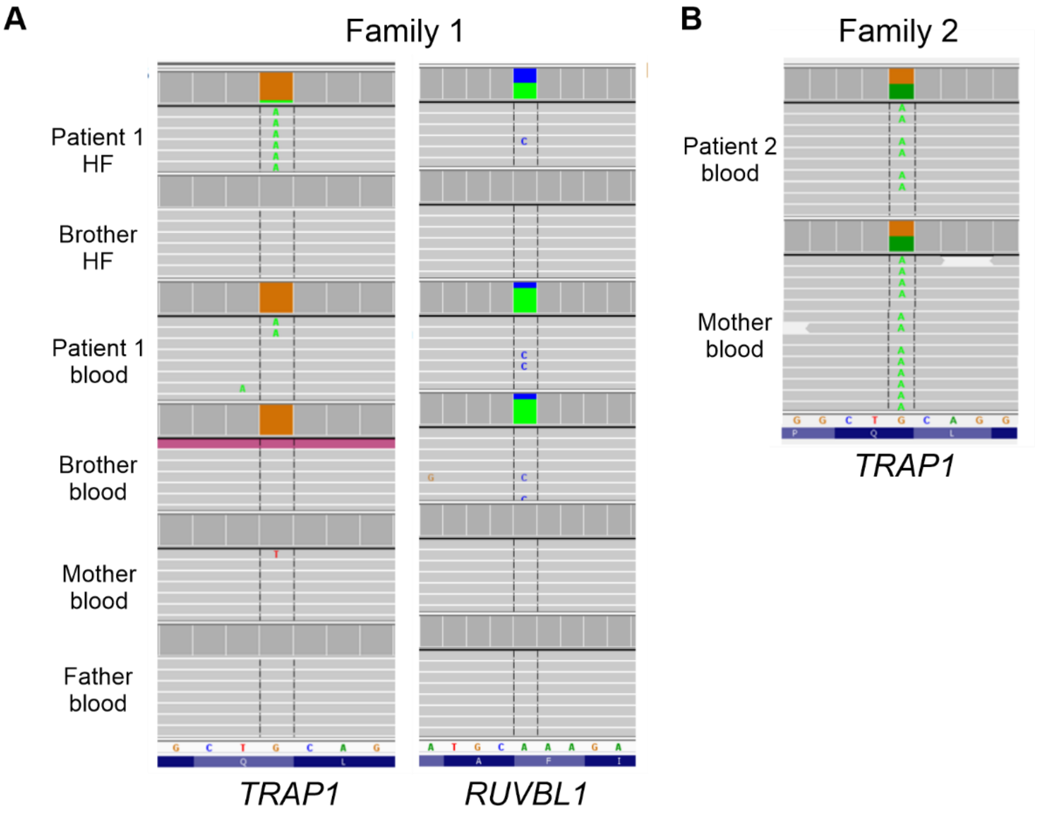


**Fig. S1; related to Figure 1**

**NGS-based deep amplicon sequencing of postzygotic *TRAP1* and *RUVBL1* variants identified by whole exome sequencing in a pair of ASD-discordant MZTs and the *TRAP1* p.Q639* variant from a replication cohort individual.**

(**A**) Variants *TRAP1* p.Q639* and *RUVBL1* p.F329L were verified in the ASD-affected twin and his ASD-unaffected twin brother in DNA samples purified from hair follicles (HF) and blood; parental analysis was done on blood samples only. In the ASD affected twin the VAF of *TRAP1* p.Q639* in HF DNA sample was 8% (genomic position coverage 13368x), and in the blood DNA sample, the VAF was 2% (genomic position coverage 27384x). In the unaffected brother HF DNA the variant was not present (genomic position coverage 7779x), while in the blood, the VAF was 2% (genomic position coverage 31972x). In the parent samples, only the wild-type sequence was identified (coverage 26714x for mother and 28222x for father). In the ASD affected twin, the VAF of RUVBL1 p.F329L in HF DNA was 48% (coverage 54610x) and in the blood sample the VAF was 22% (coverage 49383x). In the unaffected brother's HF DNA only the wild-type sequence was identified (coverage 52237x), while in blood the VAF was 22% (coverage 40726x). In the parents, only the wild-type sequence was identified (coverage 57498x for mother and 51788x for father). (**B**) Verification of the heterozygous *TRAP1* p.Q639* variant in an ASD patient from the replication cohort revealed inheritance from a ASD-unaffected mother (VAF 50% in both proband and mother); DNA from the proband's father was not available for testing. Deep amplicon sequencing results were viewed with the Integrative Genomics Viewer (IGV) tool.

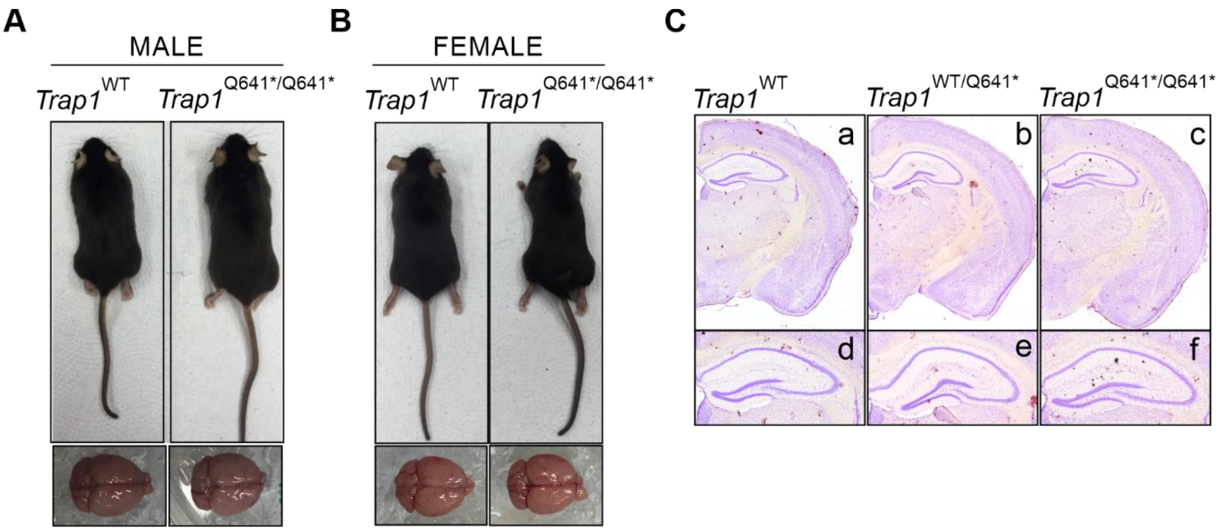


**Fig. S2; related to Figure 2**

**Phenotypic characterization of adult *Trap1*$^{WT}$ and *Trap1*$^{Q641*/Q641*}$ mice.**

**(A, B)** Photos of *Trap1*$^{WT}$ and *Trap1*$^{Q641*/Q641*}$ male and female mice at 12 weeks of age. No clear aberrant phenotype was observed in *Trap1*$^{Q641*/Q641*}$ mice. No gross anatomical differences in the brains of *Trap1*$^{Q641*}$ mice were observed. **(C)** Nissl-stained coronal sections of brains from wild type **(a, d)** heterozygous **(b, e)** and homozygous mutant **(c, f)** mice. No gross neuroanatomical differences were observed.

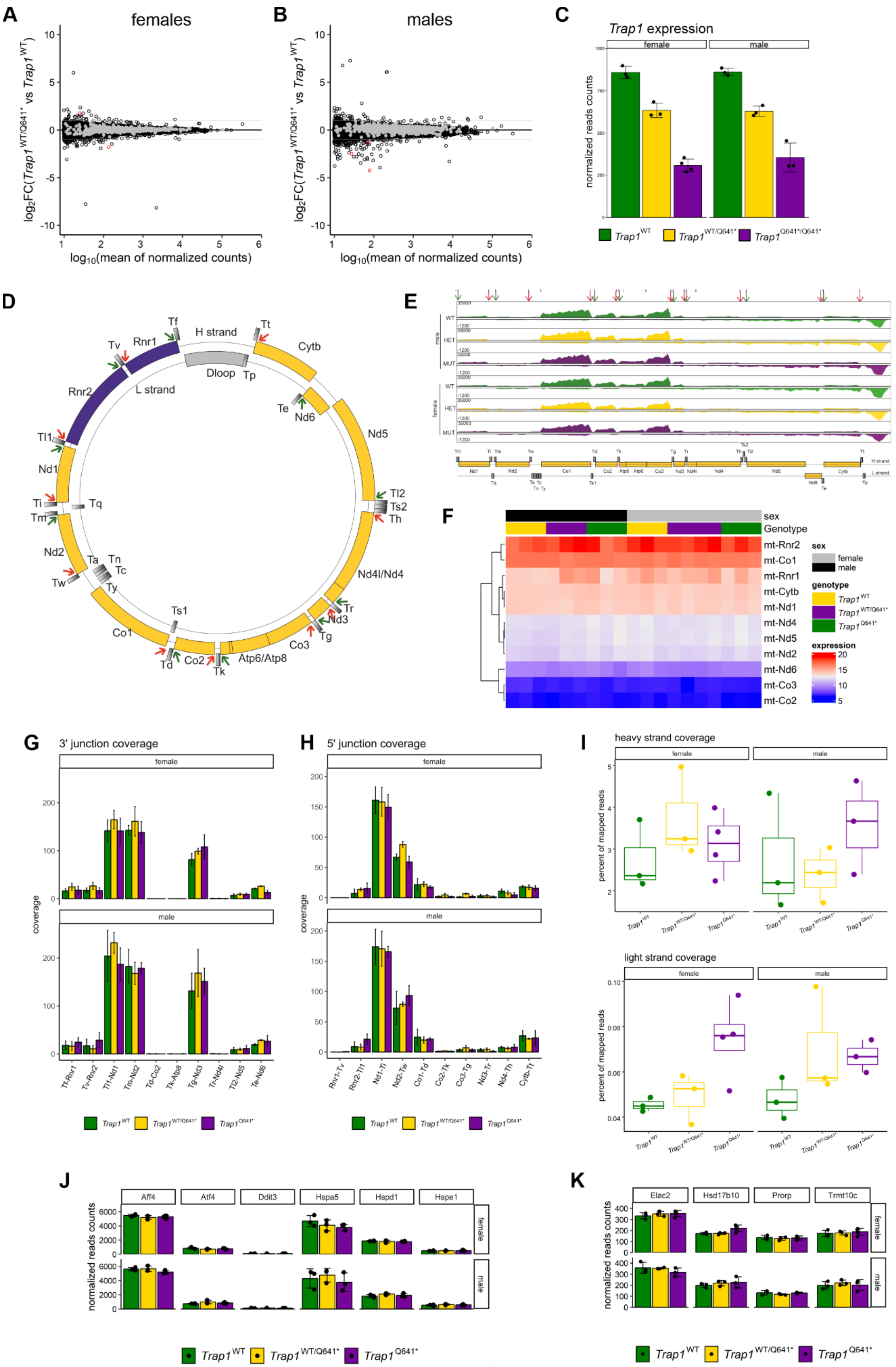
A
females
B
males
log2FC(Trap1 WT/Q641* vs Trap1 WT)
log10(mean of normalized counts)
C
Trap1 expression
female
male
normalized reads counts
Trap1 WT
Trap1 WT/Q641*
Trap1 Q641*/Q641*
D
H strand
L strand
Dloop
Rnr1
Rnr2
Cytb
Nd1
Nd2
Nd5
Nd6
Nd4l/Nd4
Nd3
Co1
Co2
Co3
Atp6/Atp8
E
male
female
WT
HET
MUT
H strand
L strand
F
sex
Genotype
mt-Rnr2
mt-Co1
mt-Rnr1
mt-Cytb
mt-Nd1
mt-Nd4
mt-Nd5
mt-Nd2
mt-Nd6
mt-Co3
mt-Co2
sex
female
male
genotype
Trap1 WT
Trap1 WT/Q641*
Trap1 Q641*
expression
G
3' junction coverage
H
5' junction coverage
coverage
I
heavy strand coverage
light strand coverage
percent of mapped reads
J
Aff4
Atf4
Ddit3
Hspa5
Hspd1
Hspe1
K
Elac2
Hsd17b10
Prorp
Trmt10c
normalized reads counts

#### Fig. S3; related to Figure 2

**RNA Sequencing analysis of the hippocampi of *Trap1*$^{WT}$, *Trap1*$^{WT/Q641*}$ and *Trap1*$^{Q641*/Q641*}$ mice**

**(A, B)** MA plots representing the global differential gene expression in RNA-Seq analysis of the hippocampi of *Trap1*$^{WT/Q641*}$ and *Trap1*$^{WT}$ in female (A) and male (B) mice. The x axis indicates the mean of normalized reads and the y axis indicates the log2 fold change of expression when comparing *Trap1*$^{WT/Q641*}$ to *Trap1*$^{WT}$. Black circles represent transcripts that are not differentially expressed, red circles represent transcripts significantly differentially expressed (|log2(FC)| >1, p.adj-value < 0.05), and gray circles represent genes encoding proteins with mitochondrial localization. (**C)** *Trap1* transcript levels as assessed in the bulk RNA-Seq analysis, plotted separately. **(D)**. Map of the mitochondrial genome. The H (heavy, outer circle) and L (light, inner circle) strands are shown with their corresponding genes. The green and red arrows indicate the 5` and 3` junctions, respectively, that were analyzed for pre-RNA processing dysregulation. **(E)** Visualization of alignment of RNA-Seq reads mapped to the mitochondrial genome. Screenshots of the per base coverage visualized with the IGV v2.3 viewer. **(F)** Heatmap representation of expression of genes encoded by the mitochondrial genome on transcript levels from the RNA-Seq analysis. The colors represent the vsd normalized reads. Blue represents low expression, whereas red represents high expression. **(G, H)** Analysis of mitochondrial pre-RNA processing defects based on the number of reads crossing the tRNA/mRNA gene junction. The reads were calculated by the number that map on the crossing junction assigned at 10 nt upstream and downstream of the real 1-nt junction. **(I)** The percentage contribution of mitochondrial reads in total mapped RNA-Seq reads for the H and L strands. **(J, K)** Levels of transcripts in the RNA-seq data encoding proteins involved in the unfolded protein response (J) and in mitochondrial tRNA processing (K).

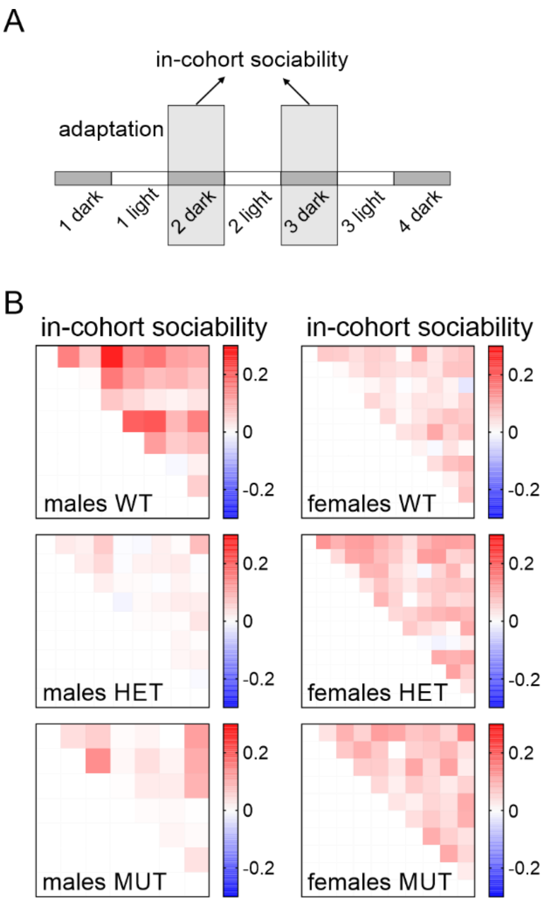


**Fig. S4; related to Figure 3**

**Eco-HAB behavioral analysis of social interactions in *Trap1*$^{WT}$, *Trap1*$^{WT/Q641*}$ and *Trap1*$^{Q641*/Q641*}$ mouse cohorts**
**(A)** Schematic depicting the experimental timeline and data used for analysis of behavioural measures. Locomotor activity was assessed in 12-hour bins and "in-cohort sociability" was calculated from the data collected during the 2$^{nd}$ and 3$^{rd}$ dark phase. **(B)** Heat maps depicting "in-cohort sociability" results. Each small square represents the "sociability" parameter for one pair of subjects, the colour scale used is *blue* to *red*, with the *blue* representing "low sociability" and *red* "high sociability". N=7-12 animals/group. Frequency distribution histograms for all pairs of animals from different cohorts are presented in main Figure 3D-E.

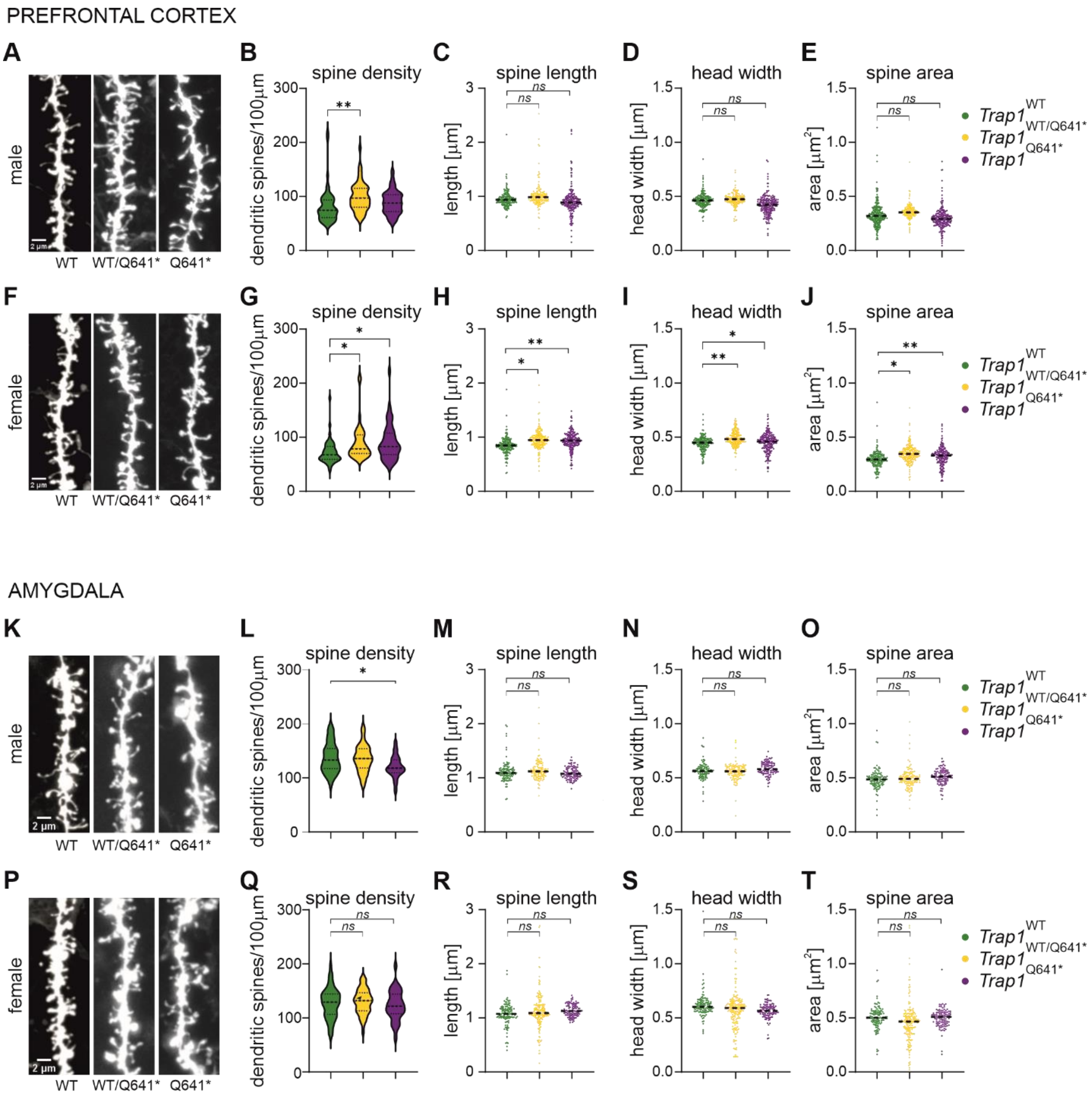


### Fig. S5; related to Figure 4

**Morphometric analysis of dendritic spines in the medial prefrontal cortex and amygdala of *Trap1*$^{WT}$, *Trap1*$^{WT/Q641*}$ and *Trap1*$^{Q641*/Q641*}$ mice**

**(A, F, K, P)** Representative images of DiI stained dendrites in the medial prefrontal cortex (mPFC) **(A, F)** and amygdala **(K, P)** of male and female *Trap1* mice. Scale bars 2 µm. **(B, G)** Mean density of dendritic spines in mPFC. Plot shows mean density/100 µm of dendrite; n=35-53 (males), n=41-44 (females) images/group; *p<0.05, **p<0.01; one way ANOVA, *post-hoc* Tukey's test. **(C-E, H-J)**. Dendritic spine morphology in *Trap1* male (C-E) and female (H-J) medial prefrontal cortex. Plots show mean value for spine length, spine head width, and spine area; n= 3727-6746 (males), 4350-5500 (females) analyzed spines/experimental group; *p<0.05, **p<0.01,***p<0.001; nested ANOVA, *post-hoc* Tukey's test. N=3-6 animals/group (males), N=3-4 animals/group (females). **(L, Q)** Mean density of dendritic spines in the amygdala. Plot shows mean density/100 µm of dendrite; n=34-65 (males), n=32-52 (females) images/group; *p<0.05, **p<0.01; one way ANOVA, *post-hoc* Tukey's test. **(M-O, R-T)** Dendritic spine morphology in *Trap1* male and female amygdala. Plots show mean value for spine length, spine head width, and spine area; n= 2496-4760 (males), 2695-4184 (females) analyzed spines/experimental group; *p<0.05, **p<0.01,***p<0.001; nested ANOVA, *post-hoc* Tukey's test. N=3-6 animals/group (males), N=3-5 animals/group (females).

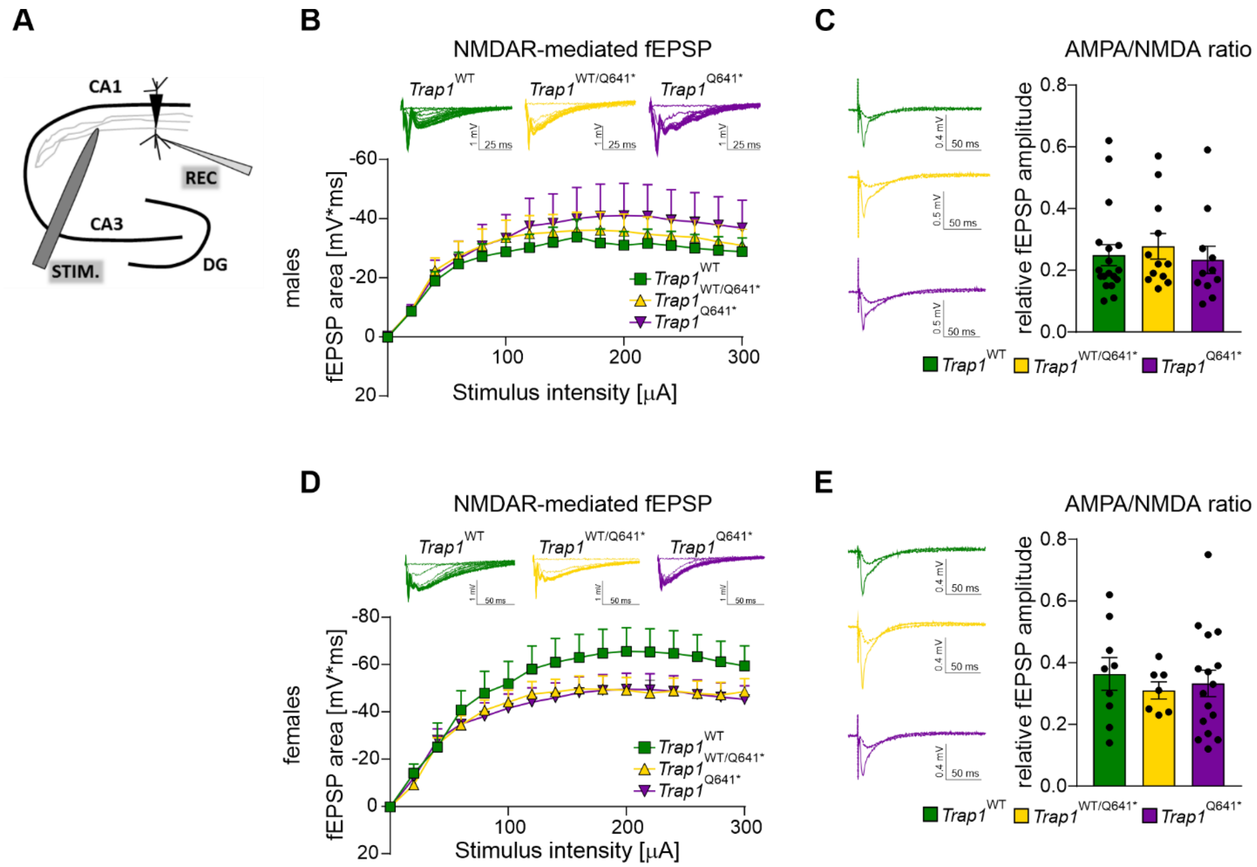


#### Fig. S6, related to Figure 4

**Electrophysiological recordings of compound AMPAR- and NMDAR-mediated fEPSPs in the CA1 hippocampal region of *Trap1* mice.**

**(A)** Schematic of the electrophysiological recording setup depicting positions of stimulating (STIM) and recording (REC) electrodes in the CA1 hippocampal region. **(B)** Averaged synaptic responses recorded in response to monotonically increasing stimuli applied to presynaptic fibers. Male *Trap1*$^{Q641*/Q641*}$ and *Trap1*$^{WT/Q641*}$ mice did not significantly differ in NMDARs-mediated synaptic responses compared to WT littermates (Kruskal-Wallis test, $p>0.05$). **(C)** Quantification of fEPSPs amplitude change following application of AMPAR antagonist DNQX (20μM). Sensitivity to DNQX and thus AMPAR/NMDAR ratio was not significantly different among male groups (One-Way ANOVA, $p>0.05$). **(D)** Averaged synaptic responses recorded in females. *Trap1*$^{Q641*/Q641*}$ and *Trap1*$^{WT/Q641*}$ groups had significantly reduced NMDARs-mediated synaptic responses compared to wild-type litter mates (Kruskal-Wallis test, $p<0.001$, Dunn's Method for multiple pairwise comparison). **(E)** Quantification of fEPSPs amplitude change following application of AMPAR antagonist DNQX (20μM). Sensitivity to DNQX was not significantly different among investigated groups (One-Way ANOVA, $p>0.05$). Insets in panels B, D show example recordings of compound fEPSPs in response to monotonically increasing stimuli (B,D) or before and after DNQX application (C,E). N=3-6 animals, n=12-25 slices (males); N=3-4 animals, n=12-17 slices (females).

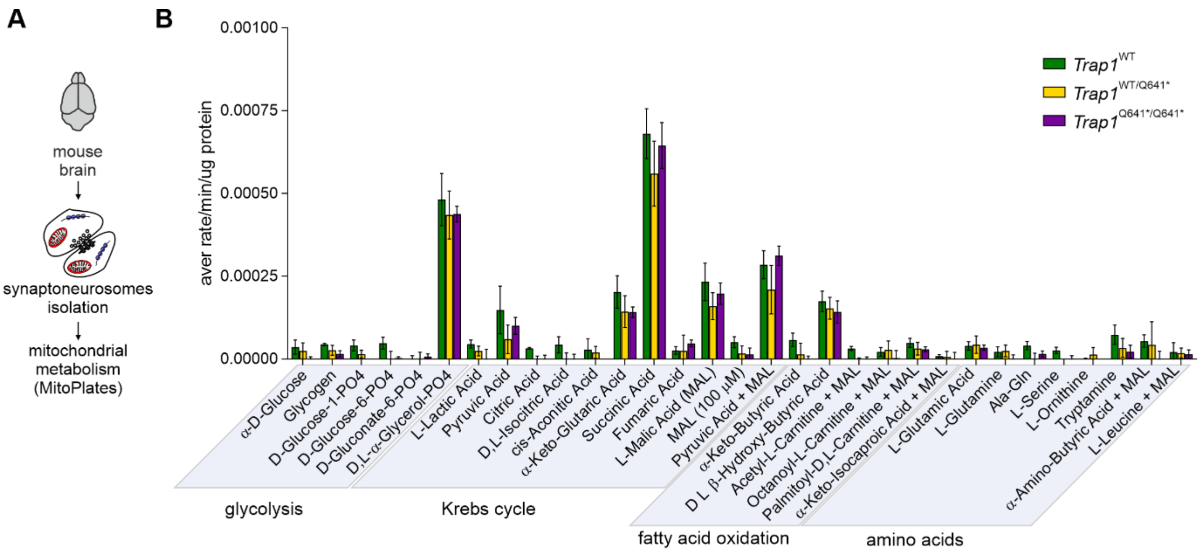


**Fig. S7, related to Figure 5**

**Functional mitochondrial phenotyping of synaptoneurosomes isolated from mouse brains of female *Trap1* mice.** The rates of production of NADH and $FADH_2$ from 31 different bioenergetic substrates, including glycolysis, TCA cycle intermediates, fatty acids and amino acids, were measured using MitoPlate™. No differences in utilization of mitochondrial energy substrates were observed. Results are presented as the average rate/min/µg of protein, +/- SEM ($n$ = 3 per genotype; two-way ANOVA, *post hoc* Sidak's multiple comparisons test).

**Table S1, related to Figure 1**
**Results of psychological evaluation of ASD-discordant MZTs**

| Patient | **MZT-6_1** | MZT-6_2 | **MZT-22_1** | MZT-22_2 | **MZT-29_1** | MZT-29_2 |
|---|---|---|---|---|---|---|
| **Age at sample collection** | 5 years old | | 7 years old | | 9 years old | |
| **Diagnosis** | **Childhood autism** | None | **Asperger's syndrome** | None | **Autism Spectrum Disorder** | None |
| **ADOS Classification** | **ASD** | Non-spectrum | **Autism** | Non-spectrum | **Autism** | Non-spectrum |
| **ADOS Communication** | **3** | 3 | **6** | 1 | **4** | 2 |
| **ADOS Social Interaction** | **8** | 0 | **12** | 2 | **12** | 1 |
| **ADOS Restricted and Repetitive Behavior** | **5** | 0 | **2** | 0 | **4** | 0 |
| **ADI-R Classification** | **Autism** | Non-spectrum | **Autism** | Non-spectrum | **Autism** | Non-spectrum |
| **ADI-R Social Interaction** | **18** | 1 | **28** | 1 | **19** | 3 |
| **ADI-R Communication** | **9** | 2 | **24** | 3 | **12** | 4 |
| **ADI-R Restricted and Repetitive Behavior** | **5** | 0 | **11** | 0 | **7** | 0 |
| **ADI-R Onset** | **5** | 2 | **4** | 2 | **4** | 3 |

bolded – ASD affected twin, ADOS - Autism Diagnostic Observation Schedule (*3*), ADI-R - Autism Diagnostic Interview – Revised (*1, 2*); highlighted columns refer to the Patient 1 and his unaffected twin brother

#### Table S2, related to Figure 1

**Clinical characteristic of 176 unrelated ASD patients from the Polish population (replication cohort).**

| Characteristics | | | |
|---|---|---|---|
| Total number of patients | 176 | | |
| *Gender* | n (%) | | |
| Males | 149 (84.7 %) | | |
| Females | 27 (15.3 %) | | |
| *Age of diagnosis* | | | |
| Range | 12 months – 9.5 years | | |
| Mean age ± SD | 3.92 ± 1.97 | | |
| *Phenotype Classification* [a] | Total | Males | Females |
| | n (%) | n (%) | n (%) |
| F84.0 | 135 (76.7) | 109 (73.2) | 26 (96.3) |
| F84.1 | 2 (1.1) | 2 (1.3) | 0 |
| F84.5 | 16 (9.1) | 16 (10.7) | 0 |
| F84.8 | 1 (0.6) | 1 (0.7) | 0 |
| F84.9 | 22 (12.5) | 21 (14.1) | 1 (3.7) |

a – phenotype classification according to the International Statistical Classification of Diseases and Related Health Problems 10th Revision (ICD-10) -World Health Organization; Version:2016 (https://icd.who.int/browse10/2016/en). F84.0 - Childhood autism, F84.1 - Atypical autism, F84.5 - Asperger syndrome, F84.8 - Other pervasive developmental disorders, F84.9 - Pervasive developmental disorder
.

**Table S3, related to Figure 1**

**Rare, functional *TRAP1* gene variants (frequency < 0.0001) identified in replication and control groups**

| Variant (GRCh37/hg19) | Effect (NM_016292.3) | ID | Number of individual with SNV | gnomAD allele freq. [a] | Number of samples in DMG database [b] | Clinical symptoms including ASD or DID [c] | Pathogenicity prediction [d] | CADD phred score [e] | Comments |
|---|---|---|---|---|---|---|---|---|---|
| **Replication group (n=176, unrelated ASD Polish patients)** | | | | | | | | | |
| **chr16:003712013-G>A** | **p.Q639*/c.1915C>T** | | **1** | **0** | **1** | **Yes** | **nd** | **46** | patient2, inherited from ASD-unaffected mother |
| chr16:003712924-A>G | p.W585R/c.1753T>C | rs749530717 | 1 | 0.00005915 | 8 | No | T | 25.5 | not present in mother, DNA from father not available for examination |
| chr16:003714426-G>A | p.S473L/c.1418C>T | rs771557989 | 1 | 0.000006570 | 0 | - | T | 23.8 | inherited from ASD-unaffected mother |
| chr16:003714454-T>C | p.I464V /c.1390A>G | rs780375051 | 1 | 0 | 0 | - | T | 22.3 | inherited from ASD-unaffected father |
| **Control group (n=100, unrelated healthy adult men from Polish population)** | | | | | | | | | |
| chr16:003708835-G>A | p.R658C/c.1972C>T | rs139636268 | 1 | 0.00002628 | 0 | - | T | 11.17 | |
| chr16:003727515-G>A | p.Q230*/c.688C>T | rs867558133 | 1 | 0.000006570 | 0 | - | nd | 12.93 | |
| chr16:003736068-C>G | p.D134H/c.400G>C | | 1 | 0 | 1 | - | T | 20.2 | both SNVs observed in one individual (*in cis*) |
| chr16:003736071-A>G | p.S133P/c.397T>C | | | 0 | | - | T | 19.77 | |
| chr16:003740966-G>A | p.R37W /c.109C>T | rs140670575 | 1 | 0.00009200 | 0 | - | T | 8.896 | |
| chr16:003767494-G>A | p.R6W/c.16C>T | | 1 | 0.00001317 | 0 | - | T | 10.75 | |

[a] - variant frequency according to gnomAD database v3.1.2 (http://gnomad.broadinstitute.org, accessed 2023-01-20); [b] – in-house database of > 10000 Polish individuals screened by WES (Department of Medical Genetics, Medical University of Warsaw); [c] – ASD=autism spectrum disorder, DID - disorders of intellectual development; [d] - missense variant's pathogenicity prediction based on MetaSVM (*17*), T (Tolerated); nd – no data, [e] - Combined Annotation-Dependent Depletion (CADD) phred score (*18*). Bolded is variant identified in ASD-twin.
